# Ongoing, rational calibration of reward-driven perceptual biases

**DOI:** 10.1101/242651

**Authors:** Yunshu Fan, Joshua I. Gold, Long Ding

## Abstract

Decision-making is often interpreted in terms of normative computations that maximize a particular reward function for stable, average behaviors. Aberrations from the reward-maximizing solutions, either across subjects or across different sessions for the same subject, are often interpreted as reflecting poor learning or physical limitations. Here we show that such aberrations may instead reflect the involvement of additional satisficing and heuristic principles. For an asymmetric-reward perceptual decision-making task, three monkeys produced adaptive biases in response to changes in reward asymmetries and perceptual sensitivity. Their choices and response times were consistent with a normative accumulate-to-bound process. However, their context-dependent adjustments to this process deviated slightly but systematically from the reward-maximizing solutions. These adjustments were instead consistent with a rational process to find satisficing solutions based on the gradient of each monkey’s reward-rate function. These results suggest new dimensions for assessing the rational and idiosyncratic aspects of flexible decision-making.

## Introduction

Normative theory has played an important role in our understanding of how the brain forms decisions. For example, many perceptual, memory, and reward-based decisions show inherent trade-offs between speed and accuracy. These trade-offs are parsimoniously captured by a class of sequential-sampling models, such as the drift-diffusion model (DDM), that are based on the accumulation of noisy evidence over time to a pre-defined threshold value, or bound (Ratcliff, 1978; Gold and Shadlen, 2002; Bogacz et al., 2006; Krajbich et al., 2010). These models have close ties to statistical decision theory, particularly the sequential probability ratio test that can, under certain assumptions, maximize expected accuracy for a given number of samples or minimize the number of samples needed for a given level of accuracy (Barnard, 1946; Wald, 1947; Wald and Wolfowitz, 1948). However, even when these models provide good descriptions of the average behavior of groups of subjects, they may not capture the substantial variability in behavior that can occur across individuals and task conditions. The goal of this study was to better understand the principles that govern this variability, in particular how these principles relate to normative theory.

We focused on a perceptual decision-making task with asymmetric rewards. For this task, both human and animal subjects tend to make decisions that are biased towards the percept associated with the larger payoff (e.g. Maddox and Bohil, 1998; Voss et al., 2004; Diederich and Busemeyer, 2006; Liston and Stone, 2008; Serences, 2008; Feng et al., 2009; Simen et al., 2009; Nomoto et al., 2010; Summerfield and Koechlin, 2010; Teichert and Ferrera, 2010; Gao et al., 2011; Leite and Ratcliff, 2011; Mulder et al., 2012; Wang et al., 2013; White and Poldrack, 2014). These biases are roughly consistent with a rational strategy to maximize a particular reward function that depends on both the speed and accuracy of the decision process, such as the reward rate per trial or per unit time (Gold and Shadlen, 2002; Bogacz et al, 2006). This strategy can be accomplished via context-dependent adjustments in a DDM-like decision process along two primary dimensions (Fig. 1A): 1) the momentary sensory evidence, via the drift rate; and 2) the decision rule, via the relative bound heights that govern how much evidence is needed for each alternative. Subjects tend to make adjustments along one or both of these dimensions to produce overall biases that are consistent with normative theory, but with substantial individual variability (Voss et al., 2004; Cicmil et al., 2015; Bogacz et al., 2006; Simen et al., 2009; Summerfield and Koechlin, 2010; Leite and Ratcliff, 2011; Mulder et al., 2012; Goldfarb et al., 2014).

**Figure 1.**
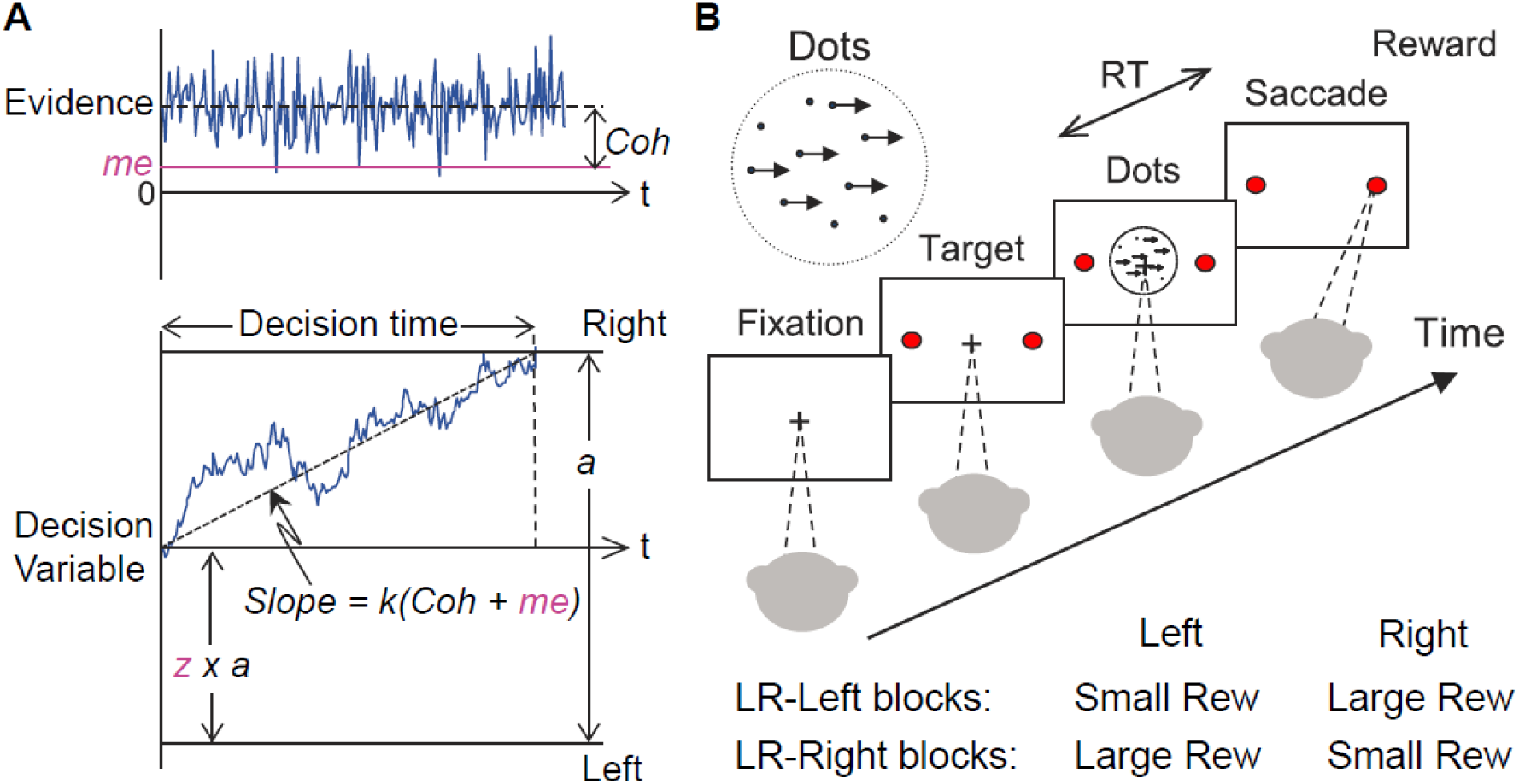
Theoretical framework and task design. A, Schematics of the drift-diffusion model (DDM). Motion evidence is modeled as samples from a unit-variance Gaussian distribution (mean: signed coherence, *Coh*). Effective evidence is modeled as the sum of motion evidence and an internal momentary-evidence bias (*me*). The decision variable starts at value a×*z*, where *z* governs decision-rule bias, and accumulates effective evidence over time with a proportional scaling factor (*k*). A decision is made when the decision variable reaches either bound. Reaction time (RT) is assumed to be the sum of the decision time and a saccade-specific non-decision time. B, Reaction-time (RT) random-dot visual motion direction discrimination task with asymmetric rewards. A monkey makes a saccade decision based on the perceived global motion of a random-dot kinematogram. Reward is delivered on correct trials and with a magnitude that depends on reward context. Two reward contexts (LR-Left and LR-Right) were alternated in blocks of trials with signaled block changes. Motion directions and strengths were randomly interleaved within blocks.

To better understand the principles that govern these kinds of idiosyncratic behavioral patterns, we trained three monkeys to perform a response-time (RT), asymmetric-reward decision task with mixed perceptual uncertainty (Fig. 1B). Like human subjects, the monkeys showed robust decision biases toward the large-reward option. These biases were sensitive to not just the reward asymmetry, as has been shown previously, but also to experience-dependent changes in perceptual sensitivity. These biases were consistent with adjustments to both the momentary evidence and decision rule in the DDM. However, these two adjustments favored the large- and small-reward choice, respectively, leading to nearly, but not exactly, maximal reward rates. We accounted for these adjustments in terms of a satisficing, gradient-based learning model that calibrated biases to balance the relative influence of perceptual and reward-based information on the decision process. Together, the results imply the broad applicability of normative theory, including sequential-sampling and heuristic-based strategies to understand how the brain combines uncertain sensory input and internal preferences to form decisions that can vary considerably across individuals and task conditions.

## Results

We trained three monkeys to perform the asymmetric-reward random-dot motion discrimination (“dots”) task (Fig. 1B). All three monkeys were initially trained on a symmetric-reward version of the task for which they were required to make fast eye movements (saccades) in the direction congruent with the global motion of a random-dot kinematogram to receive juice reward. They then performed the asymmetric-reward versions that were the focus of this study. Specifically, in blocks of 30–50 trials, we alternated direction-reward associations between a “LR-Right” reward context (the large reward was paired with a correct rightward saccade and the small reward was paired with a correct leftward saccade) and the opposite “LR-Left” reward context. We also varied the ratio of large versus small reward magnitudes (“reward ratio”) across sessions for each monkey. Within a block, we randomly interleaved motion stimuli with different directions and motion strengths (expressed as coherence, the fraction of dots moving in the same direction). We monitored the monkey’s choice (which saccade to make) and RT (when to make the saccade) on each trial.

### The monkeys’ biases reflected changes in reward context and perceptual sensitivity

For the asymmetric-reward task, all three monkeys tended to make more choices towards the large-reward option, particularly when the sensory evidence was weak. These choice biases corresponded to horizontal shifts in the psychometric function describing the probability of making a rightward choice as a function of signed motion coherence (negative for leftward motion, positive for rightward motion; Fig. 2A). We quantified monkeys’ perceptual sensitivity and bias magnitude by fitting a logistic function to monkeys’ choice behavior (example fits are shown in Supplementary Fig. 1). These measurements differed in detail for the three monkeys. For example, bias magnitude tended to be smallest in monkey F. Moreover, bias magnitude tended to decrease with session number for all three monkeys, although this tendency was statistically significant only for monkey C after accounting for co-variations with reward rate (Fig. 2B, middle). Additionally, each monkey showed steady increases in perceptual sensitivity (steepness of the psychometric function), which initially dropped relative to values from the symmetric-reward task then tended to increase with more experience with asymmetric rewards (Fig. 2B, top; *H_0_*: partial Spearman’s *ρ* of sensitivity versus session index after accounting for session-specific reward ratios = 0, *p*<0.01 in all cases, except LR-Left for monkey C, for which *p*=0.57). These improvements did not involve systematic changes in lapse rates, which remained near zero across sessions (Fig. 2B, bottom), implying that the monkeys knew how to perform the task. However, these increases in sensitivity did correspond to systematic decreases in choice biases in some, but not all, cases (3 monkeys x 2 reward contexts; Fig. 2C), which is expected for an optimal process that maximizes reward per trial (RTrial; Supplementary Fig. 2C). Compared to these optimal biases, the monkey’s biases tended to vary over sessions and overshoot for monkeys F and C (Fig. 2D).

**Figure 2.**
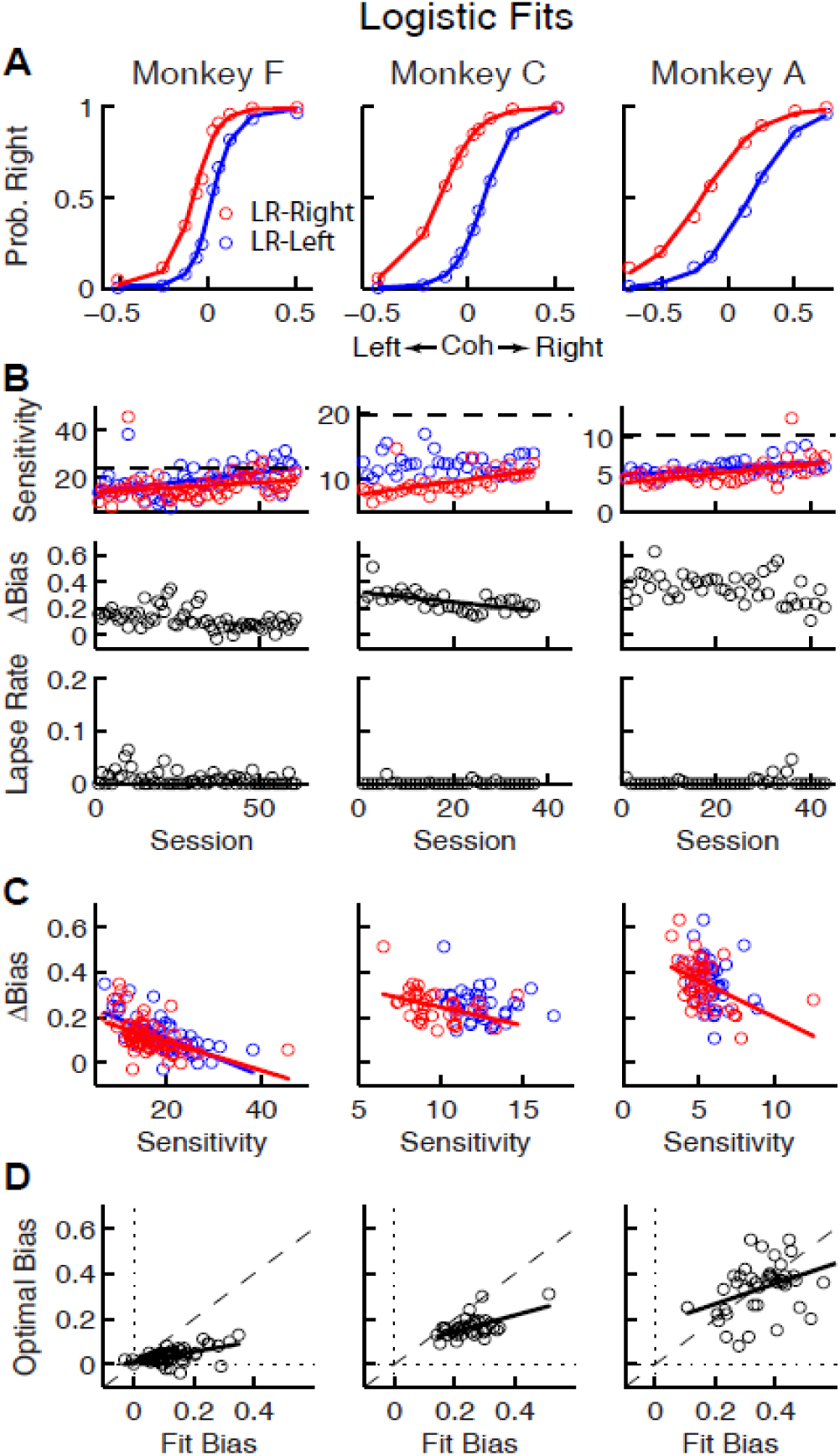
Relationships between sensitivity and bias from logistic fits to choice data. A, For each monkey, the probability of making a rightward choice is plotted as a function of signed coherence (–/+ indicate left/right motion) from all sessions, separately for the two reward contexts, as indicated. Lines are logistic fits. B, Top row: Motion sensitivity (steepness of the logistic function) in each context as a function of session index (colors as in A). Solid lines indicate significant positive partial Spearman correlation after accounting for changes in reward ratio across sessions (*p*<0.05). Black dashed lines indicate each monkey’s motion sensitivity in the same task with equal rewards before training on this asymmetric reward task. Middle row: ΔBias (horizontal shift between the two psychometric functions in the two reward contexts at chance level) as a function of session index. Solid line indicates significant negative partial Spearman correlation after accounting for changes in reward ratio across sessions (*p*<0.05). Bottom row: Lapse rate as a function of session index (median=0 for all three monkeys). C, ΔBias as a function of motion sensitivity for each reward context (colors as in A). Solid line indicates a significant negative partial Spearman correlation after accounting for changes in reward ratio across sessions (*p*<0.05). D, Optimal versus fitted Δbias. Optimal Δbias was computed as the difference in the horizontal shift in the psychometric functions in each reward context that would have resulted in the maximum reward per trial, given each monkey’s fitted motion sensitivity and experienced values of reward ratio and coherences from each session (see Fig. S2a). Solid lines indicate significant negative Spearman correlation (*p*<0.01). Partial Spearman correlation after accounting for changes in reward ratio across sessions are also significant for money F and C (p<0.01).

**Figure S1.**
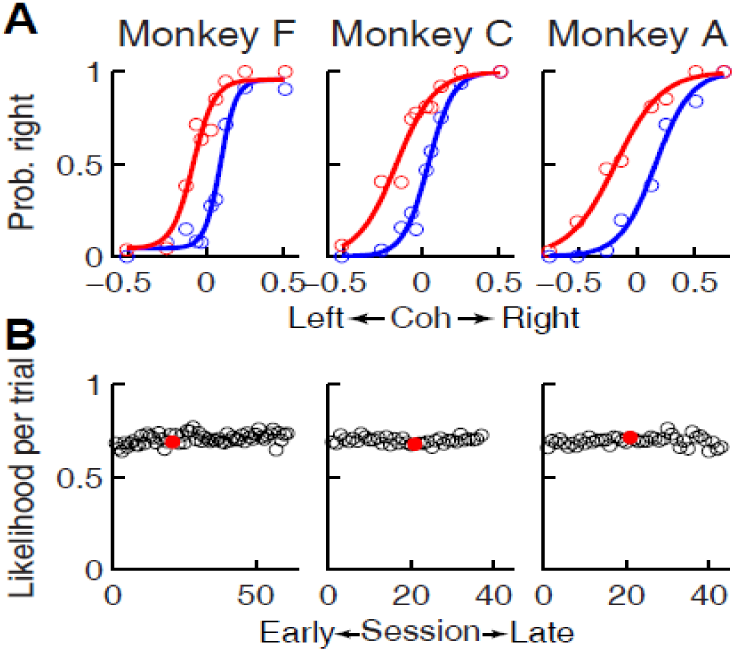
Logistic regression fits. A, Choice data from example sessions for each of the three monkeys, plotted as in Fig. 2A. Lines: fits with logistic regressions. B, Scatterplots of mean likelihood per trial, assuming binomial errors. Red circles indicate the sessions shown in A.

**Figure S2.**
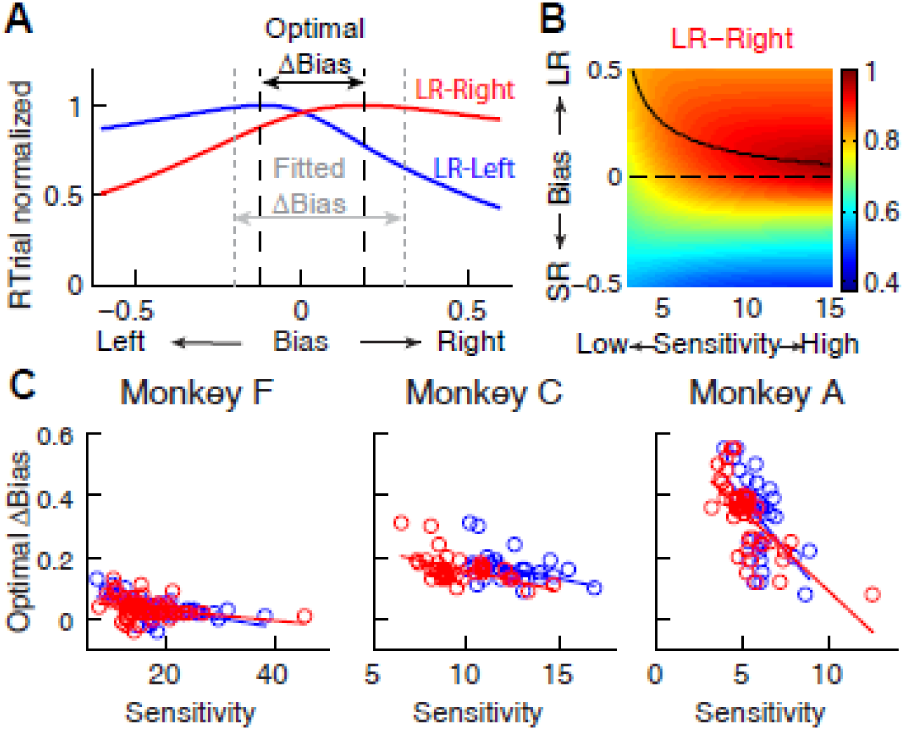
A, Identification of the optimal Δbias for an example session using logistic fits. For each reward context (blue for LR-Left and red for LR-Right), RTrial was computed as a function of bias values sampled uniformly over a broad range, given the session-specific sensitivities, lapse rate, coherences and large:small reward ratio. The optimal Δbias was defined as the difference between the bias values with the maximal RTrial for the two reward contexts. The fitted Δbias was defined as the difference between the fitted bias values for the two reward contexts. B, The optimal bias decreases with increasing sensitivity. The example heatmap shows normalized RTrial as a function of sensitivity and bias values in the LR-Right blocks, assuming the same coherence levels as used for the monkeys and a large:small reward ratio of 2.3. The black curve indicates the optimal bias values for a given sensitivity value. C, Scatterplots of optimal Δbiases obtained via the procedure described above as a function of sensitivity for each of the two reward contexts. Same format as Figure 3B. Solid lines indicate significant partial Spearman correlation after accounting for changes in reward ratio across sessions (*p*<0.05). Note that the scatterplots of the monkeys’ Δbiases and sensitivities in Fig. 2C also show negative correlations, similar to this pattern.

To better understand the computational principles that governed these idiosyncratic biases, while also taking into account systematic relationships between the choice and RT data, we fit single-trial RT data (i.e., we modeled full RT distributions, not just mean RTs) from individual sessions to a DDM. We used a hierarchical-DDM (HDDM) method that assumes that parameters from individual sessions are samples from a group distribution (Wiecki et al., 2013). The HDDM had six parameters for each reward context. Four were from a basic DDM (Fig. 1A): *a,* the total bound height, representing the distance between the two choice bounds; *k*, a scaling factor that converts sensory evidence (motion strength and direction) to the drift rate; and *t_0_* and *t_1_*, non-decision times for leftward and rightward choices, respectively. The additional two parameters provided biases that differed in terms of their effects on the full RT distributions (Supplementary Fig. S3): *me*, which is additional momentary evidence that is added to the motion evidence at each accumulating step and has asymmetric effects on the two choices and on correct versus error trials (positive values favor the rightward choice); and *z*, which determines the decision rules for the two choices and tends to have asymmetric effects on the two choices but not on correct versus error trials (values >0.5 favor the rightward choice). The HDDM fitting results are shown in Fig. 3, and summaries of best-fitting parameters and goodness-of-fit metrics are provided in Supplementary Table 1. A DDM variant with collapsing bounds provided qualitatively similar results as the HDDM (Supplementary Fig. 4). Thus, subsequent analyses use the model with fixed bounds, unless otherwise noted.

**Figure S3.**
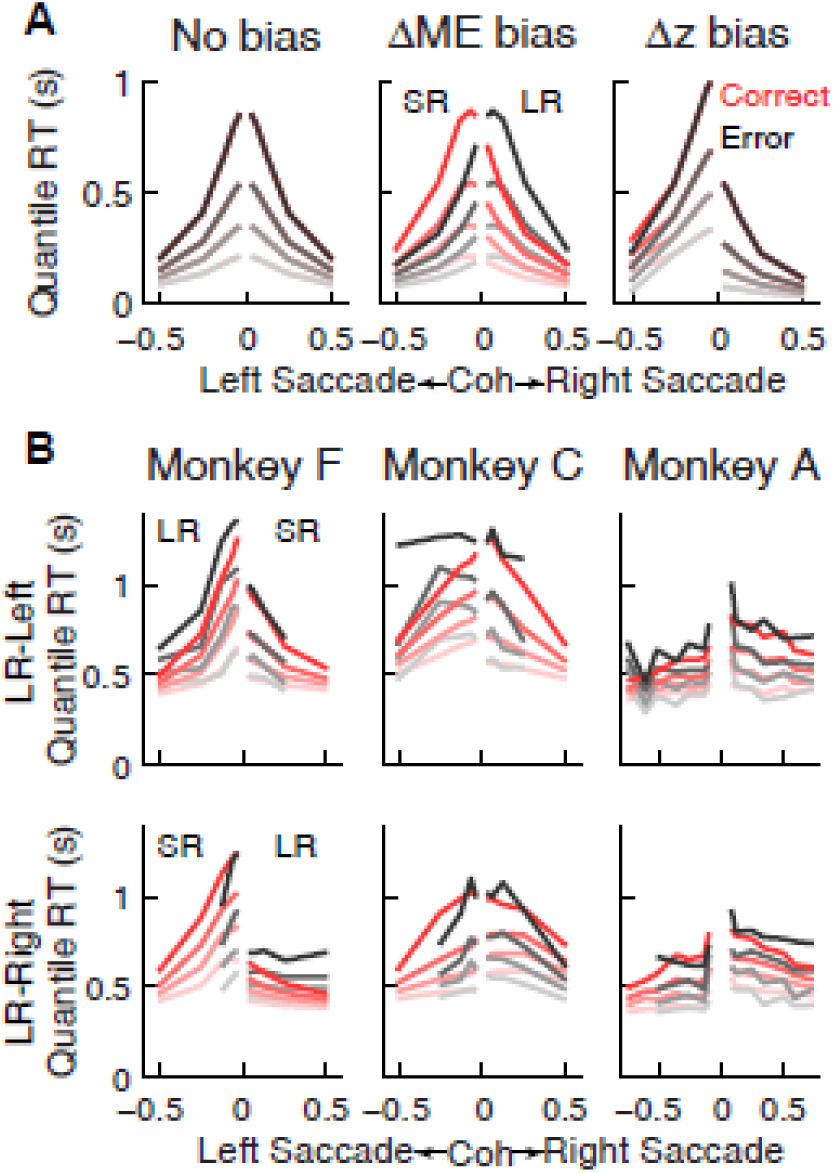
Qualitative comparison between the monkeys’ RT distribution and DDM predictions. A, RT distributions as predicted by a DDM with no bias in decision rule (*z*) or momentary evidence (*me*; left), with *me*>0 (middle), and with *z*>0.5 (right). RT distributions are shown separately for correct (red) and error (black) trials and using values corresponding to 20th, 40th, 60th, and 80th percentiles. Positive/negative coh values indicate rightward/leftward saccades. The values of *me* and *z* were chosen to induce similar choice biases (∼0.075 in coherence units). Note that the *me* bias induces large asymmetries in RT both between the two choices and between correct and error trials, whereas the *z* bias induces a large asymmetry in RT for the two choices, but with little asymmetry between correct and error trials. B, The monkeys’ mean RTs for four quantiles for the LR-Right (*top*) and LR-Left (*bottom*) reward contexts, respectively (same convention as in A). Note the presence of substantial asymmetries between correct and error trials for all three monkeys.

**Figure 3.**
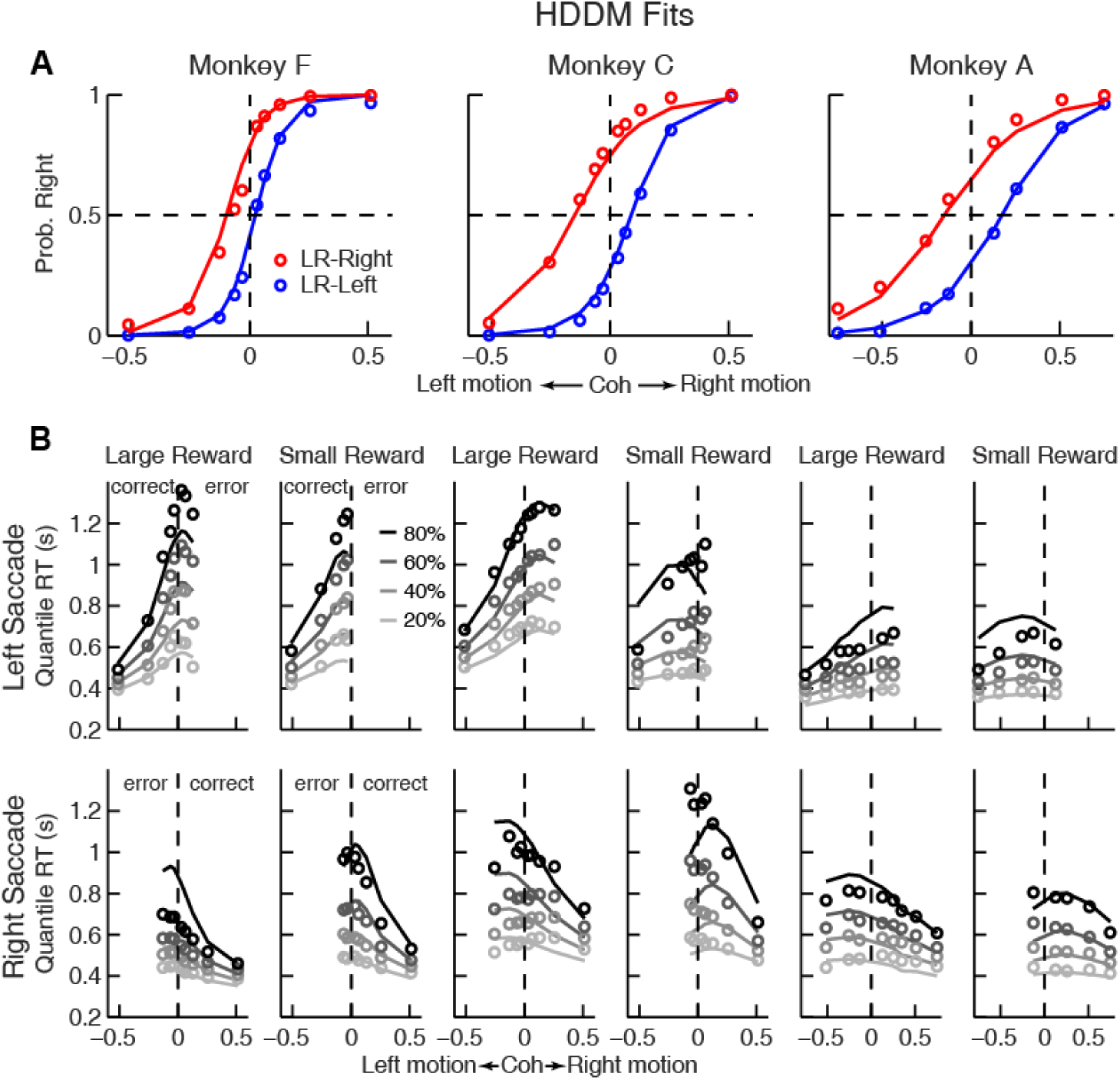
Comparison of choice and RT data to HDDM fits with both momentary-evidence (*me*) and decision-rule (*z*) biases. A, Psychometric data (points as in Fig. 2A) shown with predictions based on HDDM fits to both choice and RT data. B, RT data (circles) and HDDM-predicted RT distributions (lines). Both sets of RT data were plotted as values corresponding to the 20^th^, 40^th^, 60^th^, and 80^th^ percentiles of the full distribution. Top row: Trials in which monkey chose the left target. Bottom row: Trials in which monkeys chose the right target. Columns correspond to each monkey (as in A), divided into choices in the large- (left column) or small- (right column) reward direction (correct/error choices are as indicated in the left-most columns; note that no reward was given on error trials). The HDDM-predicted RT distributions were generated with 50 runs of simulations, each run using the number of trials per condition (motion direction × coherence × reward context × session) matched to experimental data and using the best-fitting HDDM parameters for that monkey.

**Supplementary Table 1.** Best-fitting parameters of HDDM.

|  | Monkey F (26079 trials) |  |  |  | Monkey C (37161 trials) |  |  |  | Monkey A (21089 trials) |  |  |  |
| --- | --- | --- | --- | --- | --- | --- | --- | --- | --- | --- | --- | --- |
|  | LR-Left |  | LR-Right |  | LR-Left |  | LR-Right |  | LR-Left |  | LR-Right |  |
|  | Mean | Std | Mean | Std | Mean | Std | Mean | Std | Mean | Std | Mean | Std |
| $a$ | 1.67 | 0.16 | 1.43 | 0.12 | 1.77 | 0.09 | 1.53 | 0.13 | 1.33 | 0.13 | 1.36 | 0.09 |
| $k$ | 10.22 | 1.87 | 9.91 | 2.11 | 6.58 | 0.51 | 5.08 | 0.92 | 4.04 | 0.33 | 3.45 | 0.46 |
| $t_1$ | 0.31 | 0.03 | 0.29 | 0.03 | 0.35 | 0.04 | 0.33 | 0.05 | 0.29 | 0.04 | 0.27 | 0.04 |
| $t_0$ | 0.28 | 0.04 | 0.31 | 0.05 | 0.33 | 0.04 | 0.31 | 0.03 | 0.21 | 0.08 | 0.26 | 0.04 |
| $z$ | 0.60 | 0.03 | 0.57 | 0.04 | 0.62 | 0.03 | 0.40 | 0.04 | 0.57 | 0.06 | 0.39 | 0.04 |
| $me$ | -0.06 | 0.04 | 0.08 | 0.05 | -0.14 | 0.04 | 0.21 | 0.06 | -0.22 | 0.05 | 0.27 | 0.09 |

**Figure S4.**
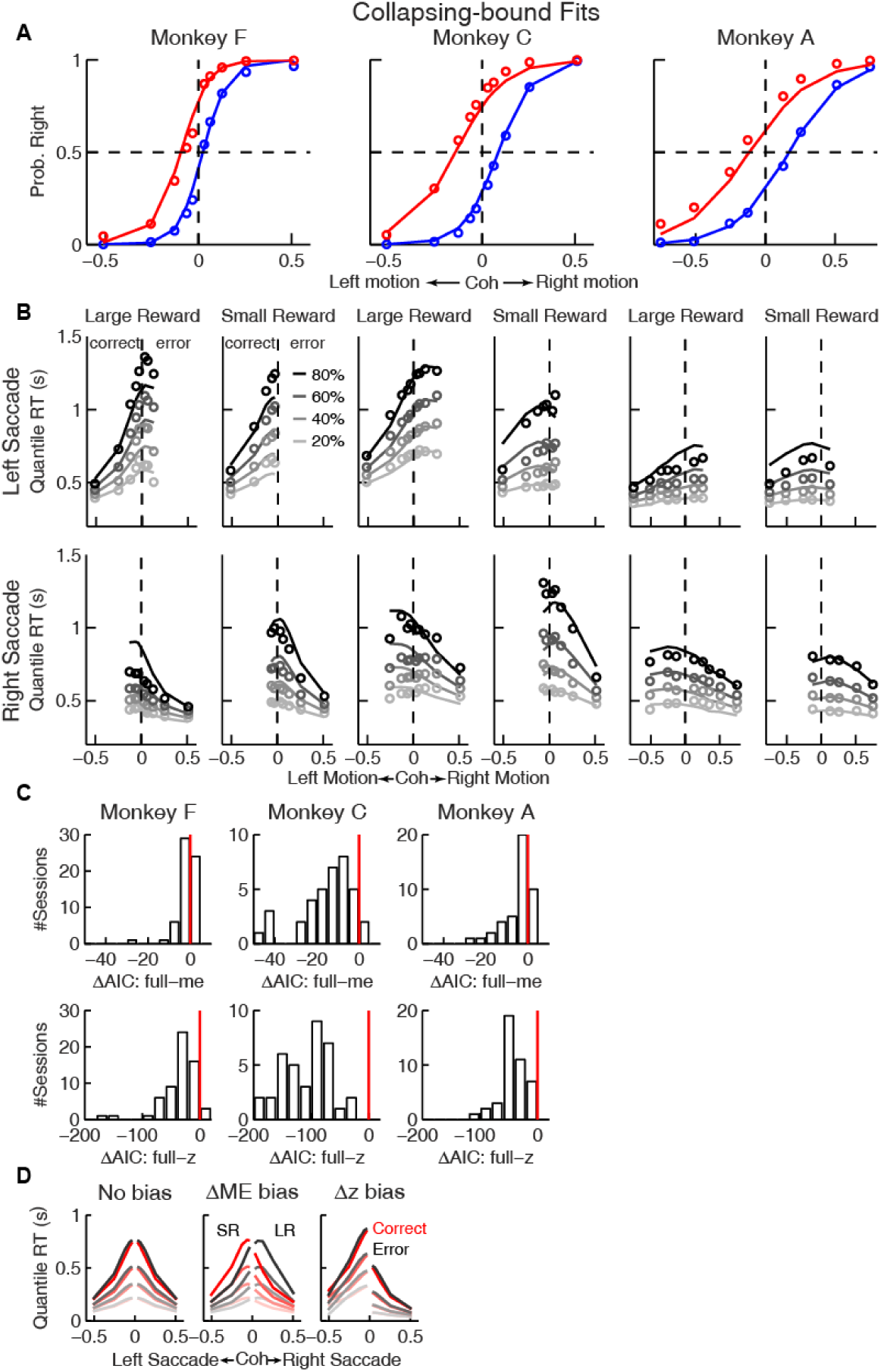
Fits to a DDM with collapsing bounds. A, B, A DDM with collapsing bounds and both momentary evidence (*me*) and decision rule (*z*) biases fit to each monkey’s RT data. Same format as Fig. 3. C, The model that included both *me* and *z* adjustments (“full”) had smaller Akaike Information Criterion (AIC) values than reduced models (“*me*” or “*z*” only) across sessions. Note also the different ranges of ΔAIC for the full–me and full–z comparisons. The mean ΔAICfull-me and ΔAICfull-z values are significantly different from zero (Wilcoxon signed rank test, *p*=0.0007 for Monkey F’s full–me comparison and *p*<0.0001 for all others). D, RT distributions as predicted by the DDM with collapsing bounds, using no bias in *z* or *me* (*left*), *me*>0 (*middle*), or *z*>0.5 (*right*). Same format as Fig. S3A.

**Supplementary Table 2.** The difference in deviance information criterion (ΔDIC) between the full model (i.e., the model that includes both *me* and *z*) and either reduced model (*me-*only or *z-*only), for experimental data and data simulated using each reduced model. Negative values favor the full model. Positive values favor the reduced models. Note that the ΔDIC for the experimental data deviated substantially - toward the negative direction - from the ΔDIC for the simulated data.

|  | Experimental Data |  |  |  | Simu: *me* model |  | Simu: *z* model |  |
| --- | --- | --- | --- | --- | --- | --- | --- | --- |
| | ΔDIC: full-*me* | | ΔDIC: full-*z* | | ΔDIC: full-*me* | | ΔDIC: full-*z* | |
|  | Mean | Std | Mean | Std | Mean | Std | Mean | Std |
| Monkey F | -124.6 | 2.7 | -2560.4 | 3.9 | 3.1 | 8.7 | 0.2 | 10.5 |
| Monkey C | -1700.4 | 2.4 | -6937.9 | 0.9 | 17.5 | 10.1 | 1.8 | 1.2 |
| Monkey A | -793.6 | 2.7 | -2225.7 | 3.1 | 25.4 | 8.0 | 1.2 | 3.1 |

The DDM fits provided a parsimonious account of both the choice and RT data. Consistent with the results from the logistic analyses, the HDDM analyses showed that the monkeys made systematic improvements in psychometric sensitivity (*H_0_*: partial Spearman’s *ρ* of sensitivity versus session index after accounting for session-specific reward ratios=0, *p*<0.01 in all cases except *p*=0.06 for LR-Left for monkey A). Moreover, there was a negative correlation between psychometric sensitivity and choice bias (*H_0_*: partial Spearman’s *ρ* of sensitivity versus total bias after accounting for session-specific reward ratios=0, *p*<0.001 in all cases). These fits ascribed the choice biases to changes in both the momentary evidence (*me*) and the decision rule (*z*) of the decision process, as opposed to either parameter alone (Supplementary Table 2). These fits also indicated context-dependent differences in non-decision times, which were smaller for all large-reward choices for all three monkeys except in LR-Right context for monkey C and A (*t-* test, *p*<0.05).

### The monkeys’ bias adjustments were adaptive with respect to optimal reward-rate functions

To try to identify common principles that governed these monkey- and context-dependent decision biases, we analyzed behavior with respect to optimal benchmarks based on certain reward-rate functions. We focused on reward per unit time (RR) and per trial (RTrial), which for this task are optimized in a DDM framework by adjusting momentary-evidence (*me*) and decision-rule (*z*) biases, such that both favor the large-reward choice. However, the magnitudes of these optimal adjustments depend on other task parameters (*a, k*, *t_0_,* and *t_1_*, non-bias parameters from the DDM, plus the ratio of the two reward sizes and inter-trial intervals) that can vary from session to session. Thus, to determine the optimal adjustments, we performed DDM simulations with the fitted HDDM parameters from each session, using different combinations of *me* and *z* values (Fig. 4A). As reported previously (Bogacz et al., 2006; Simen et al., 2009), when the large reward was paired with the leftward choice, the optimal strategy used *z*<0.5 and *me*<0 (Fig. 4B, top panels, purple and orange circles for RR and RTrial, respectively). Conversely, when the larger reward was paired with the rightward choice, the optimal strategy used *z*>0.5 and *me*>0 (Fig. 4B, bottom panels).

**Figure 4.**
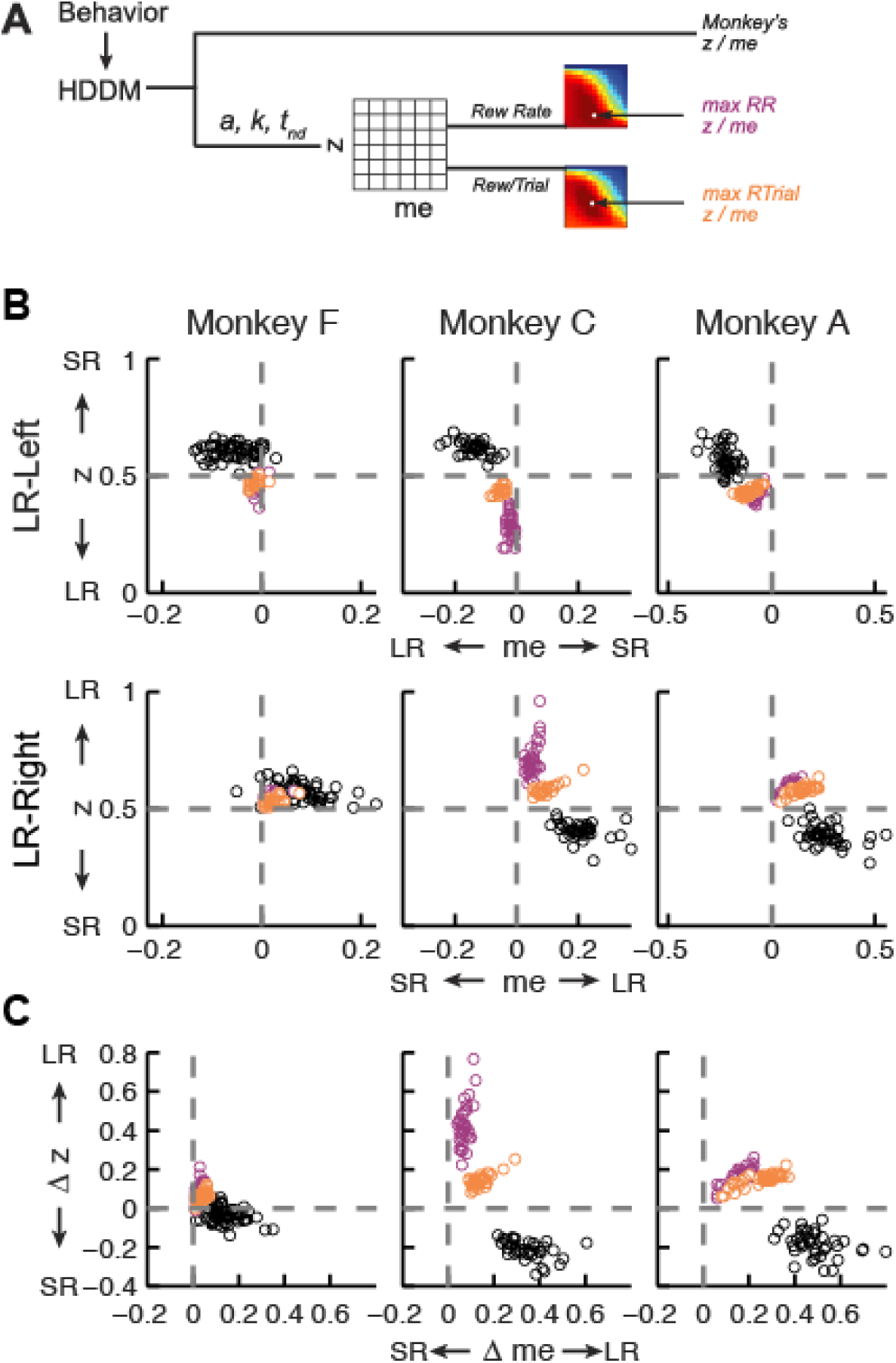
**Actual versus optimal adjustments of momentary-evidence (*me*) and decision-rule (*z*) biases.** A, Schematic of the comparison procedure. Choice and RT data from the two reward contexts in a given session were fitted separately using the HDDM. These context- and session-specific best-fitting *me* and *z* values are plotted as the monkey’s data (black circles in B and C). Optimal values were determined by taking the best-fitting values of parameters *a*, *k*, and non-decision times from the HDDM fits, then sampling *me* and *z* uniformly over broad ranges to predict choice and RT behavior. For each *me* and *z* combination, the predicted probability of left/right choice and RTs were used with actual task information (inter-trial interval, error timeout and reward sizes) to calculate the expected reward rate (RR) and average reward per trial (RTrial). Optimal *me/z* adjustments were then identified to maximize RR (purple) or RTrial (orange). B, Scatterplots of the monkeys’ *me/z* adjustments (black), predicted optimal adjustments for maximal RR (purple), and predicted optimal adjustments for maximal Rtrial (orange), for the two reward contexts in all sessions (each data point was from a single session). Values of *me*>0 or *z*>0.5 produce biases favoring rightward choices. C, Scatterplots of the differences in *me* (abscissa) and *z* (ordinate) between the two reward contexts for monkeys (black), for maximizing RR (purple), and for maximizing RTrial (orange). Positive Δme and Δz values produce biases favoring large-reward choices.

The monkeys’ adjustments of momentary-evidence (*me*) and decision-rule (*z*) biases showed both differences and similarities with respect to these optimal predictions (Fig. 4B, black circles; similar results were obtained using fits from a model with collapsing bounds, Supplementary Fig. 5). In the next section, we consider the differences, in particular the apparent use of shifts in *me* in the adaptive direction (i.e., favoring the large-reward choice) but of a magnitude that was larger than predicted, along with shifts in *z* that tended to be in the non-adaptive direction (i.e., favoring the small-reward choice). Here we focus on the similarities and show that the monkeys’ decision biases were adaptive with respect to the reward-rate function in four ways (RTrial provided slightly better predictions of the data and thus are presented in the main figures; results based on RR are presented in the Supplementary Figures).

**Figure S5.**
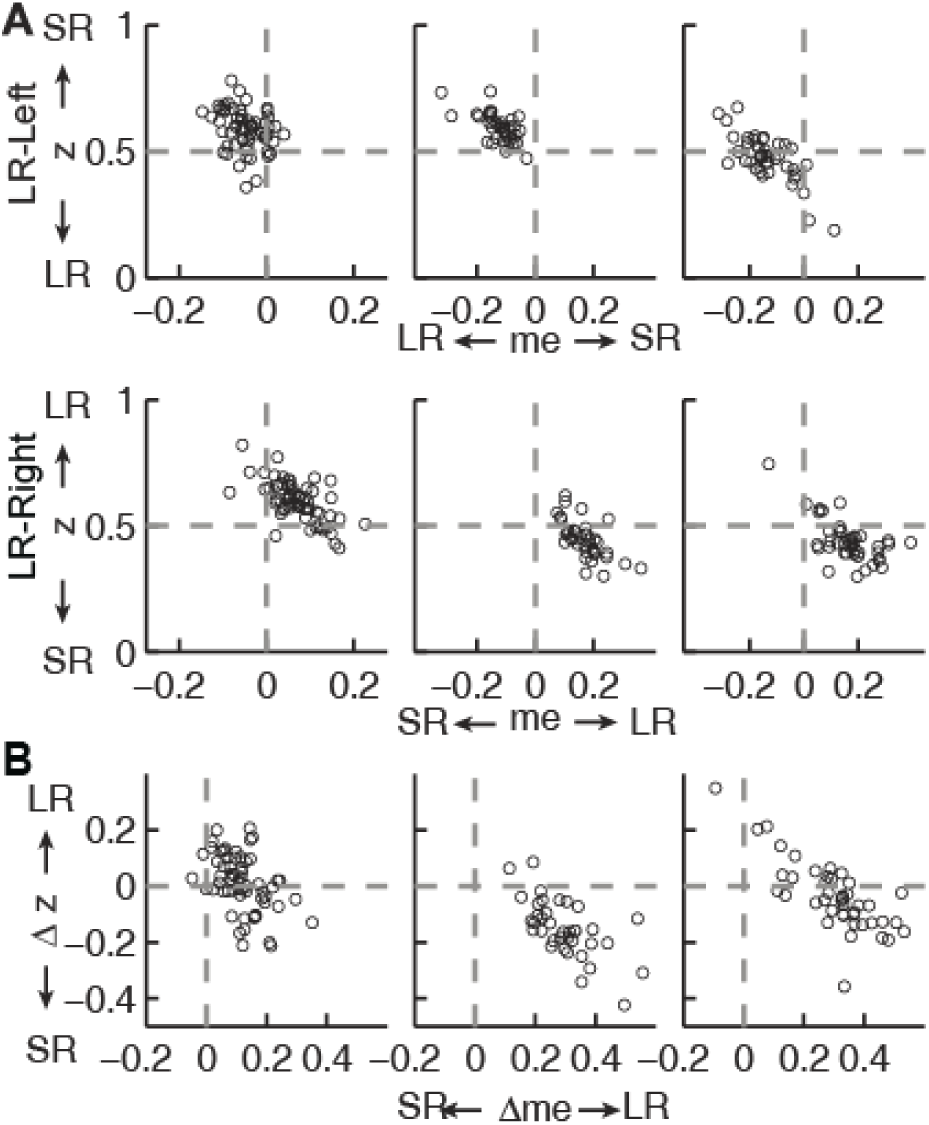
Estimates of momentary-evidence (*me*) and decision-rule (*z*) biases using the collapsing-bound DDM fits. Same format as Fig. 4B and C, except here only showing fits to the monkeys’ data. As with the model without collapsing bounds, the adjustments in *me* tended to favor the large reward but the adjustments in *z* tended to favor the small reward.

First, the best-fitting *me* and *z* values from each monkey corresponded to near-maximal reward rates (Fig. 5A). We compared the optimal values of reward per trial (RTrial_max_) to the values predicted from the monkeys’ best-fitting *me* and *z* adjustments (RTrial_predict_). Both RTrial_predict_ and RTrial_max_ depended on the same non-bias parameters in the HDDM fits that were determined per session (*a, k*, *t_0_,* and *t_1_*) and thus are directly comparable. Their ratios tended to be nearly, but slightly less than, one (mean ratio: 0.977, 0.984, and 0.982 for monkeys F, C, and A, respectively) and remained relatively constant across sessions (*H_0_*: slopes of linear regressions of these ratios versus session number=0, *p*>0.05 for all three monkeys). Similar results were also obtained using the monkeys’ realized rewards, which closely matched RTrial_predict_ (Spearman’s *ρ*=0.991, 0.979, and 0.906 for monkeys F, C, and A, respectively, *p*<0.0001 in all three cases). These results reflected the shallow plateau in the RTrial function near its peak (Fig. 5B), such that the monkeys’ actual adjustments of *me* and *z* were within the contours for 97% RTrial_max_ in most sessions (Fig. 5C; see Supplementary Fig. 6 for results using RR). Thus, the monkeys’ overall choice biases were consistent with strategies that lead to nearly optimal reward outcomes.

**Figure 5.**
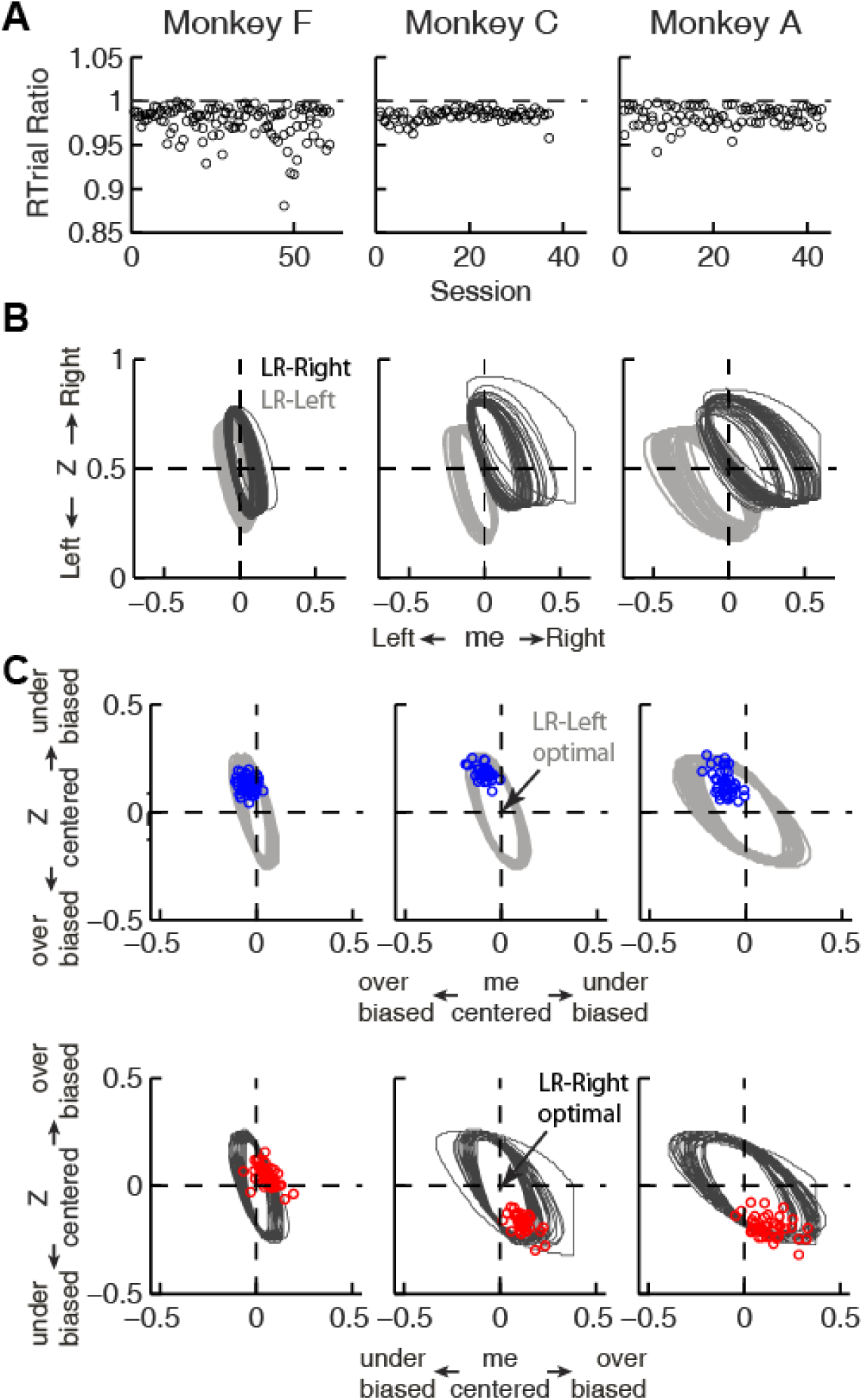
Predicted versus optimal reward per trial (RTrial). A, Scatterplots of RTrial_predict_:RTrial_max_ ratio as a function of session index. Each session was represented by two ratios, one for each reward context. Mean ratio across contexts and sessions: 0.977 for monkey F, 0.984 for monkey C, and 0.983 for monkey A. B, 97% RTrial_max_ contours for all sessions, computed using the best-fitting HDDM parameters and experienced coherences and reward ratios from each session. Light grey: LR-Left blocks; Dark grey: LR-Right blocks. C, The monkeys’ adjustments (blue in LR-Left blocks, red in LR-Right blocks) were largely within the 97% RTrial_max_ contours for all sessions and tended to cluster in the *me* over-biased, *z* under-biased quadrants (except Monkey F in the LR-Right blocks). The contours and monkeys’ adjustments are centered at the optimal adjustments for each session.

**Figure S6.**
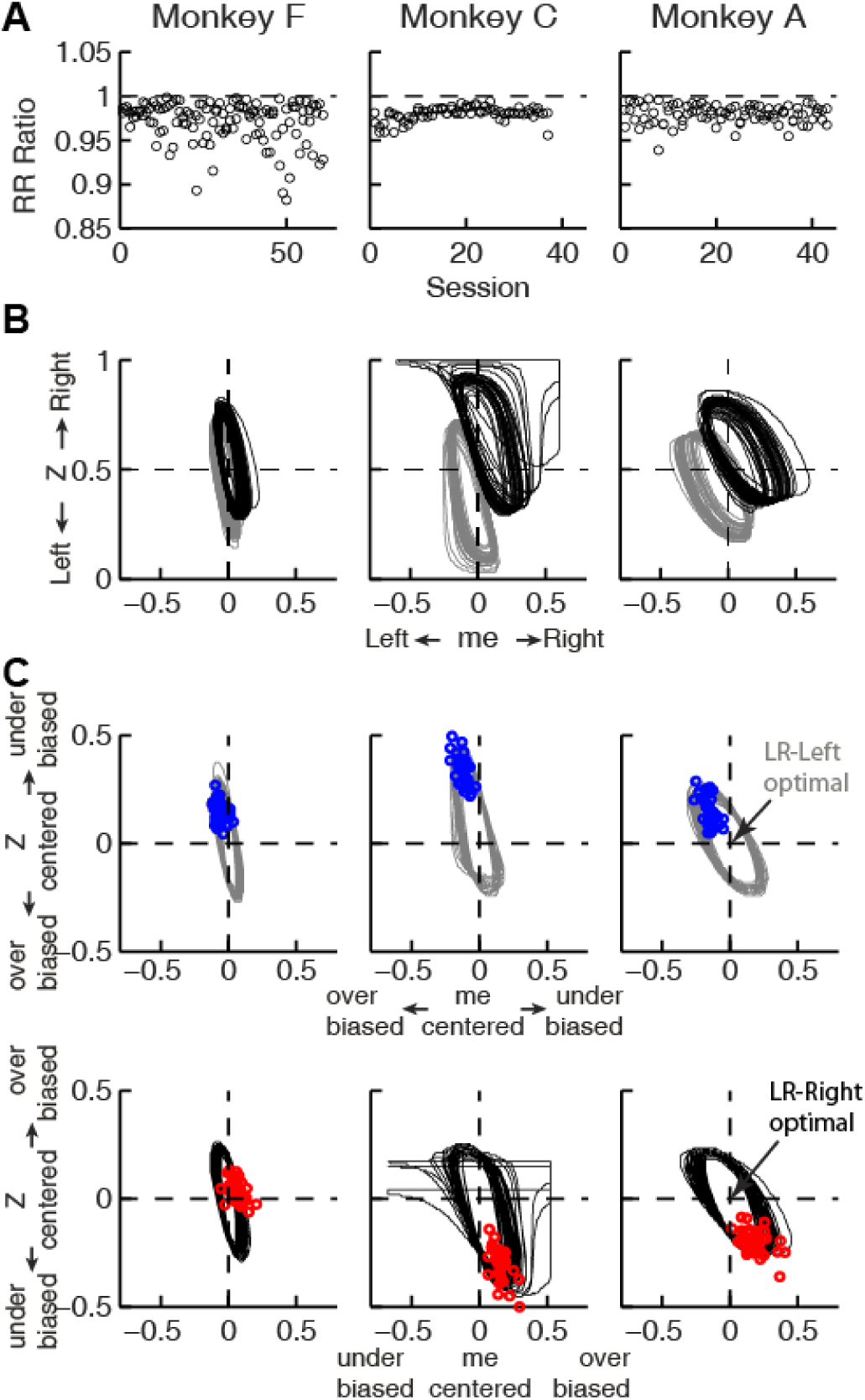
Predicted versus optimal reward rate (RR). Same format as Fig. 5. Mean RR_predict_:RR_max_ ratio across sessions=0.970 for monkey F, 0.980 for monkey C, and 0.979 for monkey A.

Second, the across-session variability of each monkey’s decision biases was predicted by idiosyncratic features of the reward functions. The reward functions were, on average, different for the two reward contexts and each of the three monkeys (Fig. 6A). These differences included the size of the near-maximal plateau (red patch), which determined the level of tolerance in RTrial for deviations from optimal adjustments in *me* and *z*. This tolerance corresponded to the session-by-session variability in each monkey’s *me* and *z* adjustments (Fig. 6B). In general, monkey F had the smallest plateaus and tended to use the narrowest range of *me* and *z* adjustments across sessions. In contrast, monkey A had the largest plateaus and tended to use the widest range of *me* and *z* adjustments (Pearson’s *ρ* between the size of the 97% RTrial contour, in pixels, and the sum of the across-session variances in each monkeys’ *me* and *z a*djustments=0.83, *p*=0.041). Analyses using the RR function produced qualitatively similar results (Supplementary Fig. 7).

**Figure 6.**
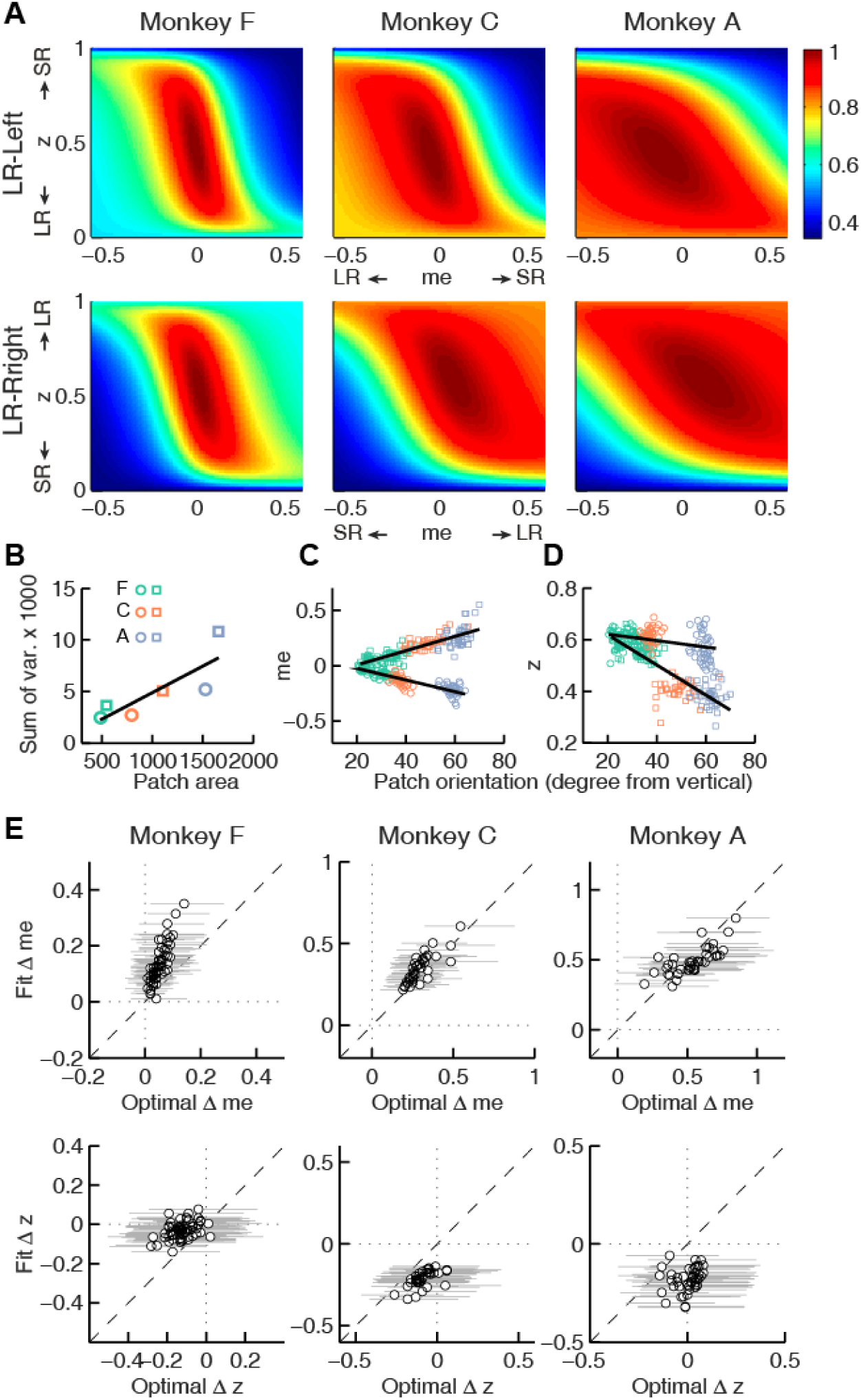
**Relationships between adjustments of momentary-evidence (*me*) and decision-rule (*z*) biases and RTrial function properties**. A, Mean RTrial as a function of *me* and *z* adjustments for the LR-Left (top) and LR-Right (bottom) blocks. RTrial was normalized to RTrialmax for each session and then averaged across sessions. B, Scatterplot of the total variance in *me* and *z* adjustments across sessions (ordinate) and the area of >97% max of the average RTrial patch (abscissa). Variance and patch areas were measured separately for the two reward blocks (circles for LR-Left blocks, squares for LR-Right blocks). C, D, The monkeys’ session- and context-specific values of *me* (C) and *z* (D) co-varied with the orientation of the >97% heatmap patch (same as the contours in Fig. 5B). Orientation is measured as the angle of the tilt from vertical. Circles: data from LR-Left block; squares: data from LR-Right block; lines: significant correlation between *me* (or *z*) and patch orientations across monkeys (*p*<0.05). Colors indicate different monkeys (see legend in B). E, Scatterplots of conditionally optimal versus fitted Δ*me* (top row) and Δ*z* (bottom row). For each reward context, the conditionally optimal *me (z)* value was identified given the monkey’s best-fitting z (*me*) values. The conditionally optimal Δ*me (*Δ*z)* was the difference between the two conditional optimal *me (z)* values for the two reward contexts. Grey lines indicate the range of conditional Δ*me* (Δ*z*) values corresponding to the 97% maximal RTrial given the monkeys’ fitted z (*me*) values.

**Figure S7.**
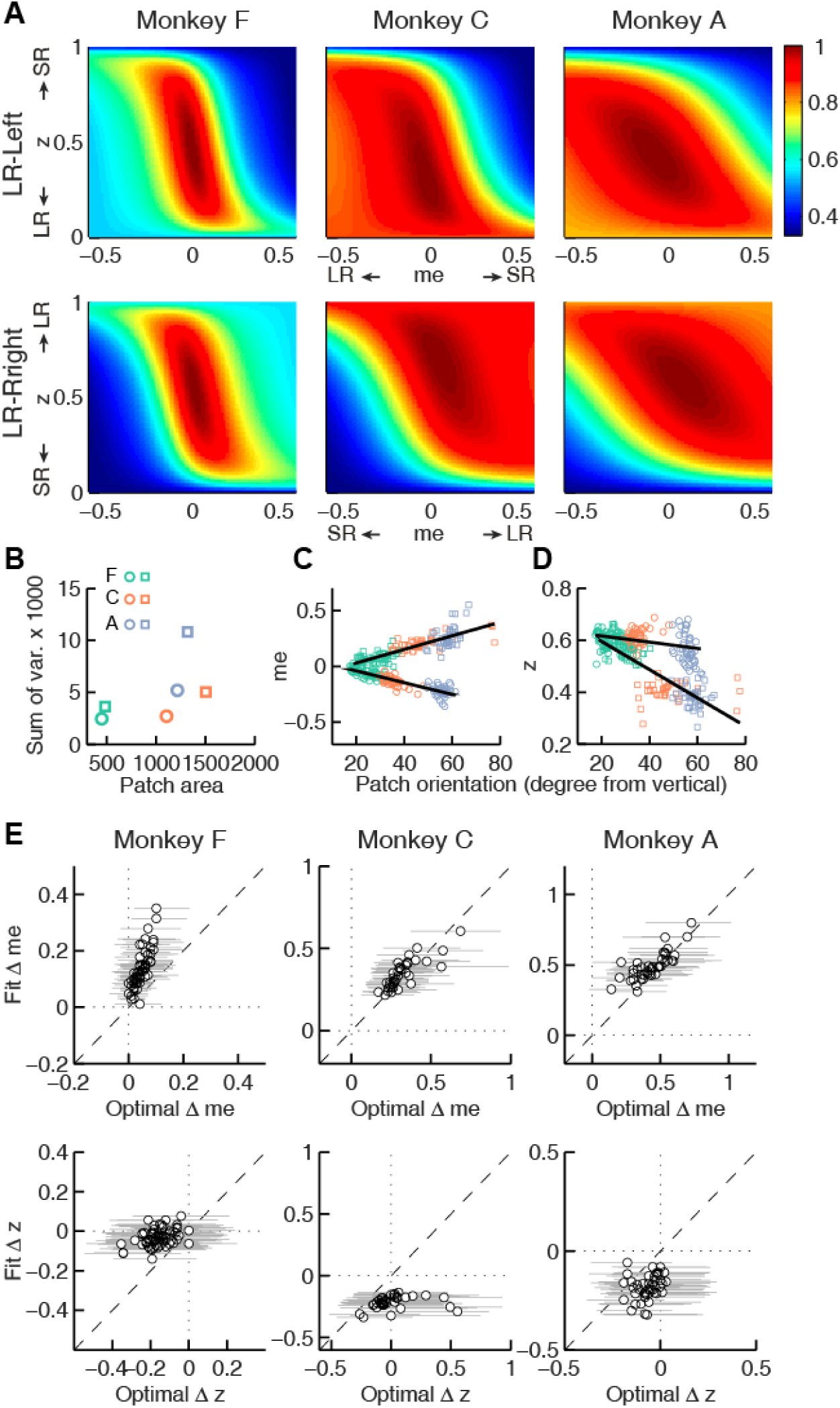
The monkeys’ momentary-evidence (*me*) and decision-rule (*z*) adjustments reflected RR function properties. Same format as Fig. 6, but using RR instead of RTrial.

**Figure S8:**
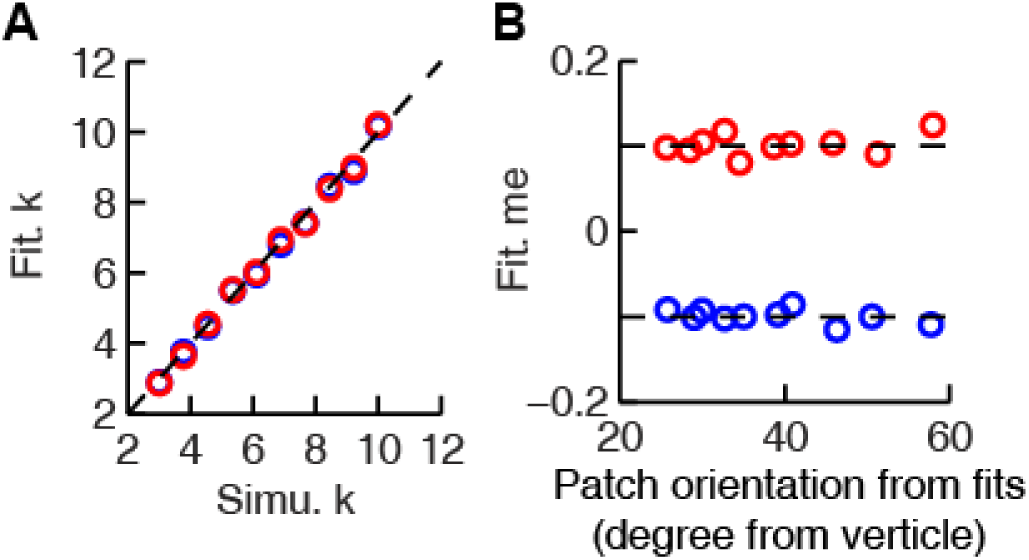
The HDDM model fitting procedure does not introduce spurious correlations between patch orientation and *me* value. Artificial sessions were simulated with fixed *me* values (±0.1 for the two reward contexts) and different *k* values. A: Recovered *k* values from HDDM fitting closely matched *k* values used for the simulations. B: Recovered *me* values from HDDM fitting closely matched *me* values used for simulation and did not correlate with RTrial patch orientation.

Third, the session-by-session adjustments in both *me* and *z* corresponded to particular features of each monkey’s context-specific reward function. The shape of this function, including the orientation of the plateau with respect to *z* and *me*, depended on the monkey’s perceptual sensitivity and the reward ratio for the given session. The monkeys’ *me* and *z* adjustments varied systematically with this orientation (Fig. 6C and D for RTrial, Supplementary Fig. 7C and D for RR). This result was not an artifact of the fitting procedure, which was able to recover appropriate, simulated bias parameter values regardless of the values of non-bias parameters that determine the shape of the reward function (Supplementary Fig. 8).

Fourth, the monkeys’ *me* and *z* adjustments were correlated with the values that would maximize RTrial, given the value of the other parameter for the given session and reward context (Fig. 6E for RTrial, Supplementary Fig. 7E for RR). These correlations were substantially weakened by shuffling the session-by-session reward functions (Supplementary Fig. 9). Together, these results suggest that all three monkeys used biases that were adaptively calibrated with respect to the reward information and perceptual sensitivity of each session.

**Figure S9.**
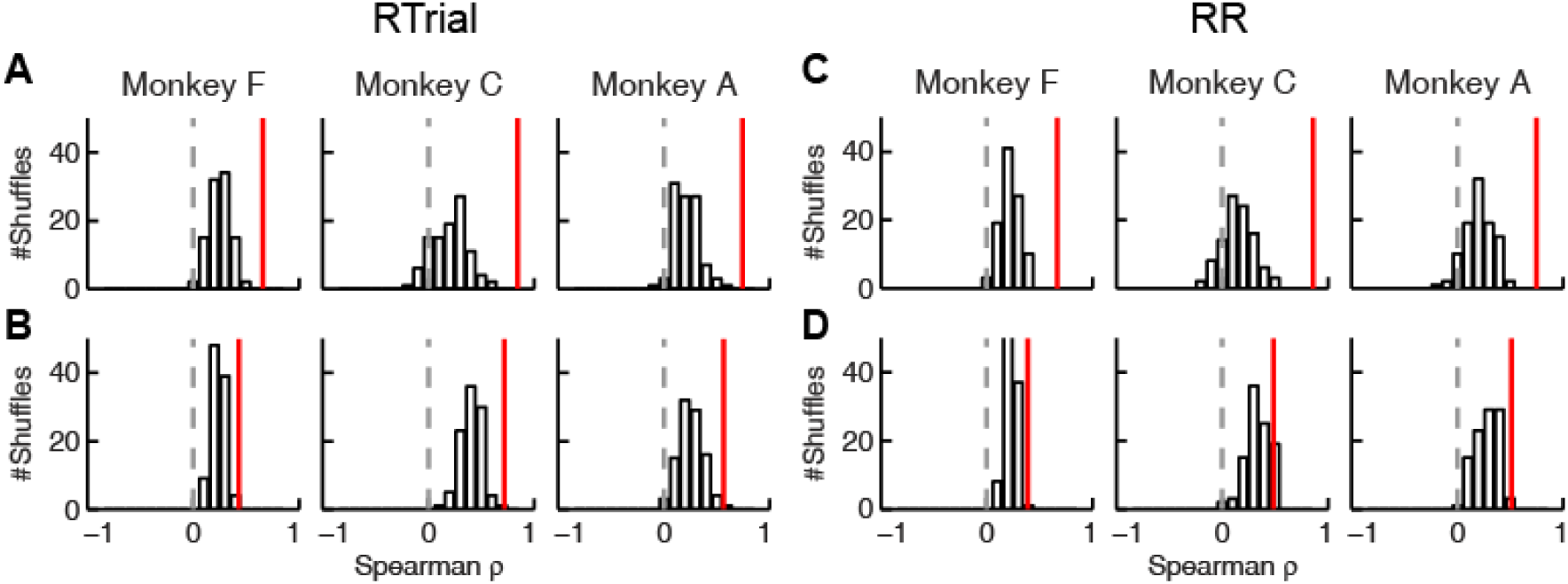
Correlation between fitted and conditionally optimal adjustments was stronger for matched sessions than for unmatched sessions. A, C: Momentary-evidence (Δ*me*) adjustments. B, D: Decision-rule (Δ*z*) adjustments. A, B: optimal values obtained with the RTrial function. C, D: optimal values obtained with the RR function. Red lines indicate the partial Spearman correlation coefficients between the fitted and optimal Δ*me* or Δ*z* (obtained in the same way as data in Fig. 6E) for matched sessions. Bars represent the histograms of partial correlation for unmatched sessions, which were obtained by 100 random shuffles of the sessions). Note that the correlation values for unmatched sessions were lower than those for matched sessions (Wilcoxson rank-sum test, *p*<0.001 for all three monkeys and both Δ*me* and Δ*z*).

### The monkeys’ adaptive adjustments were consistent with a satisficing, gradient-based learning process

Thus far, we showed that all three monkeys adjusted their decision strategies in a manner that matched many features of the optimal predictions based on their idiosyncratic, context-specific reward-rate functions. However, their biases did not match the optimal predictions exactly. Specifically, all three monkeys used shifts in *me* favoring the large-reward choice (adaptive direction) but of a magnitude that was larger than predicted, along with shifts in *z* favoring the small-reward choice (non-adaptive direction). We next consider if and how a consistent, adaptive process used by all three monkeys could lead to these idiosyncratic and not-quite optimal patterns of adjustments.

We propose that a gradient-based satisficing model can account for these patterns of adjustments. The intuition for the model is shown in Fig. 7. The lines on the RTrial heatmap represent the trajectories of a gradient-tracking procedure that adjusts *me* and *z* values to increase RTrial until reaching 97% of the maximum possible value. Gradient lines are color-coded based on how *me* and *z* values at the end points relate to the optimal *me* and *z* values. For example, consider adjusting *me* and *z* by following all of the magenta gradient lines until their endpoints. This procedure would result in *me* shifts in the adaptive direction with magnitudes larger than the optimal *me*, plus *z* shifts in the non-adaptive direction. In other words, as long as the initial *me* and *z* values fall within the area covered by the magenta lines, the positive gradient-tracking procedure would lead to a good-enough solution with over-shifted *me* and non-adaptive *z* values similar to what we found in the monkeys’ data.

**Figure 7.**
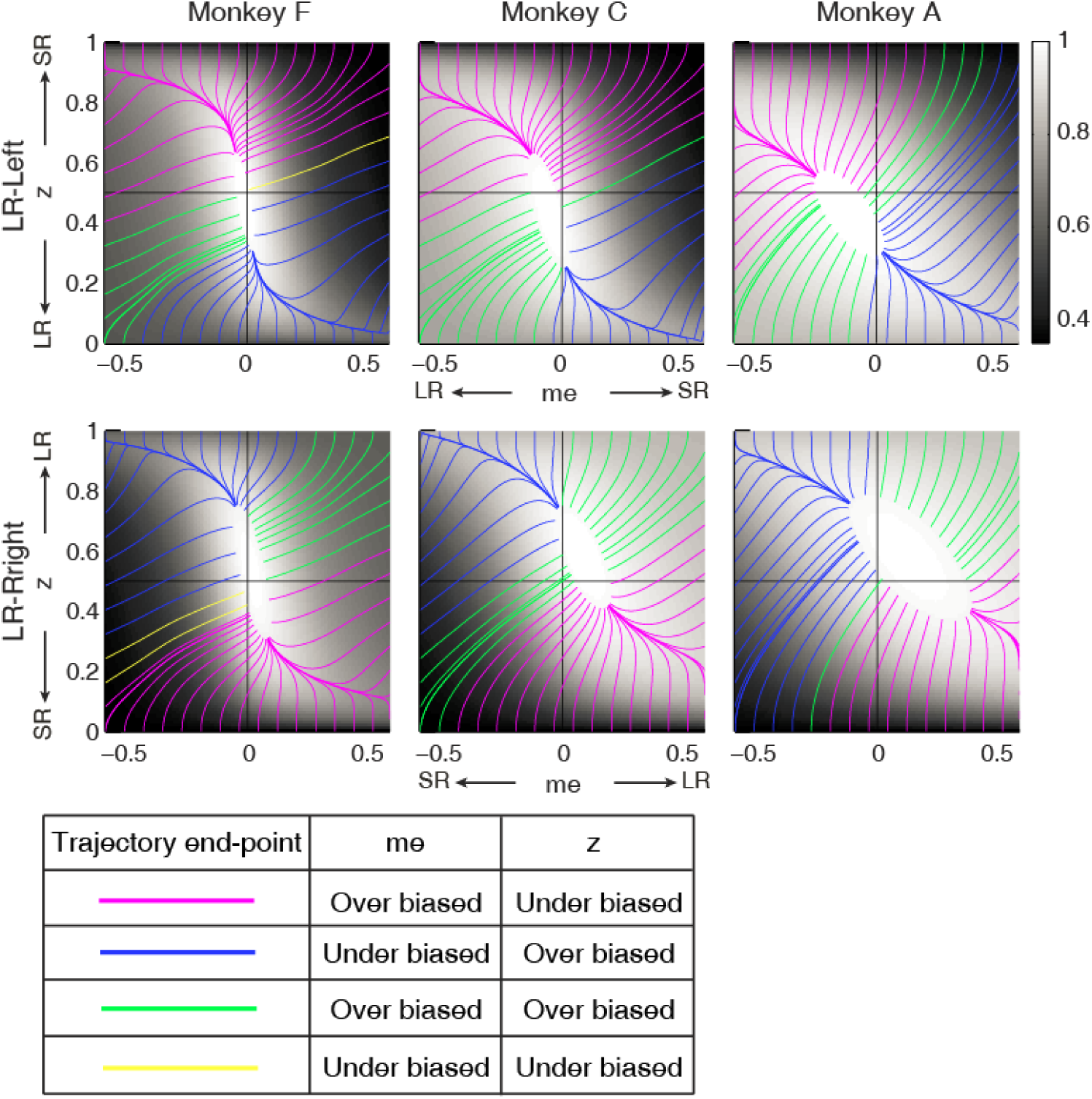
**Relationships between starting and ending values of the satisficing, reward function gradient-based updating process.** Example gradient lines of the average RTrial maps for the three monkeys are color coded based on the end point of gradient-based *me* and *z* adjustments in the following ways: 1) *me* biases to large reward whereas *z* biases to small reward (magenta); 2) *z* biases to large reward whereas *me* biases to small reward (blue); 3) *me* and *z* both bias to large reward (green), and 4) *me* and *z* both bias to small reward (yellow). The gradient lines ended on the 97% RTrialmax contours. Top row: LR-Left block; bottom row: LR-Right block.

We simulated this process with the reward function derived for each reward context, session, and monkey, using different starting points and a termination rule corresponding to achieving the estimated RTrial_predict_ from those conditions (see above). This process is illustrated for LR-Left blocks in an example session from monkey C (Fig. 8A). We estimated the unbiased *me* and *z* values as the midpoints between their values for LR-Left and LR-Right blocks (square). At this point, the RTrial gradient is larger along the *me* dimension than the *z* dimension, reflecting the tilt of the reward function. We set the initial point at baseline *z* and a very negative value of *me* (-90% of the highest coherence used in the session; overshoot in the adaptive direction) and referred to this setting as the “over-*me*” model. The *me* and *z* values were then updated in a step-wise fashion (magenta trace), according to the RTrial gradient, until the monkey’s RTrial_predict_ or better was achieved (magenta circle). The endpoint of this updating process was very close to monkey C’s actual adjustment (gray circle). For comparison, three alternative models are illustrated. The “over-*z*” model selects *z* as the initial dimension and assumes updating from the baseline *me* and over-adjusted *z* values (blue, initial *z* set as 0.1 for the LR-Left context and 0.9 for the LR-Right context). The “over-both” model assumes updating from the over-adjusted *me* and *z* values (green). The “neutral” model assumes the same updating process but from the baseline *me* and baseline *z* (black). The endpoints from these alternative models deviated considerably from the monkey’s actual adjustment.

**Figure 8.**
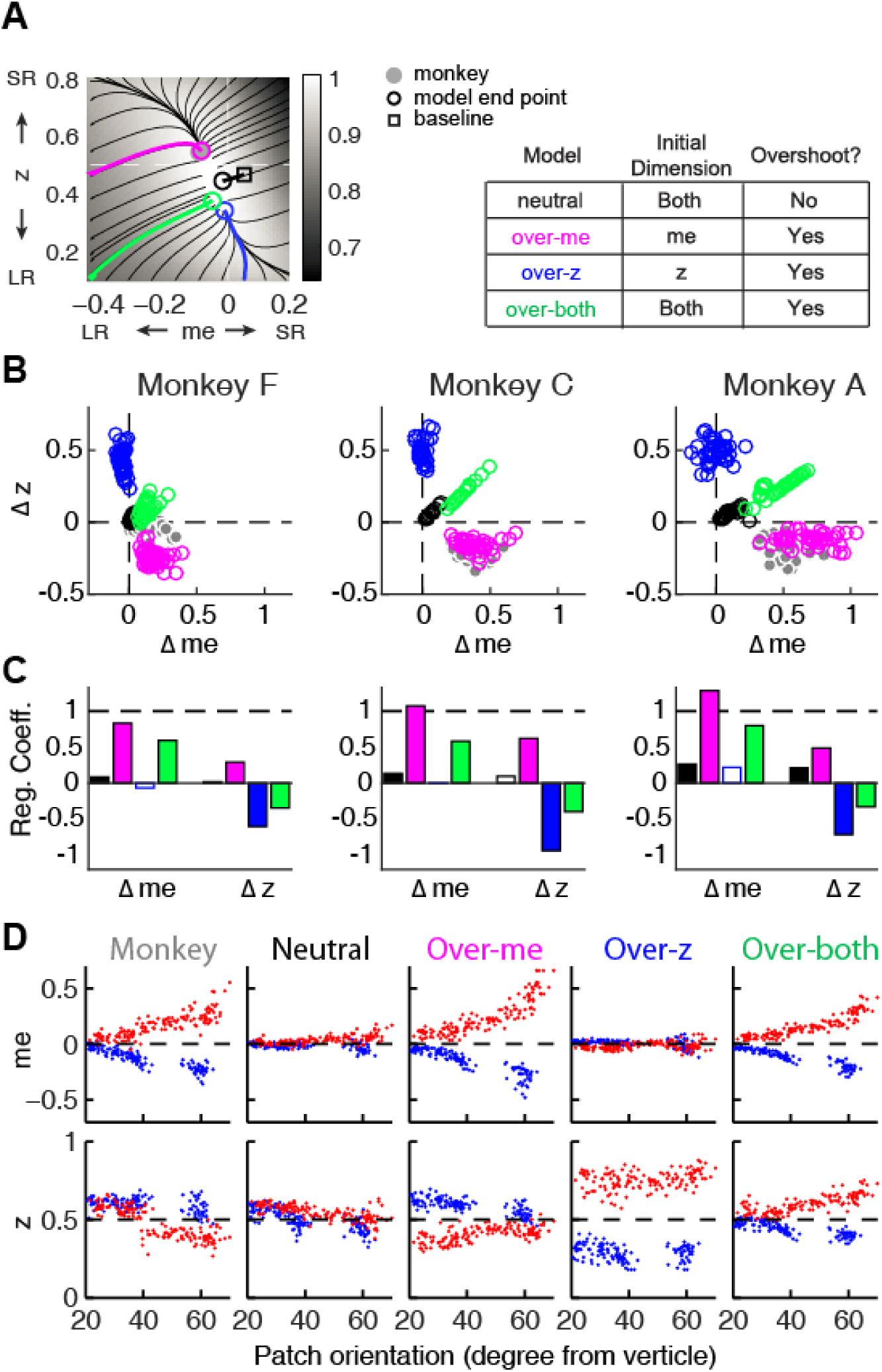
**The satisficing reward function gradient-based model.** A, Illustration of the procedure for predicting a monkey’s *me* and *z* values for a given RTrial function. For better visibility, RTrial for the LR-Left reward context in an example session is shown as a heatmap in greyscale. Gradient lines are shown as black lines. The square indicates the unbiased *me* and *z* combination (average values across the two reward contexts). The four trajectories represent gradient-based searches based on four alternative assumptions of initial values (see table on the right). Open circles indicate the end values. Grey filled circle indicates the monkey’s actual *me* and z. Note that the end points differ among the four assumptions, with the magenta circle being the closest to the monkey’s fitted *me* and *z* of that session., Scatterplots of the predicted and actual Δ*me* and Δz between reward contexts. Grey circles here are the ame as the black circles in Fig. 4C. Colors indicate model identity, as in A. B, Scatterplots of the predicted and actual Δ*me* and Δz between reward contexts. Grey circles here are the same as the black circles in Fig. 4C. Colors indicate model identity, as in A. C, Average regression coefficients between each monkey’s Δ*me* (left four bars) and Δ*z* (right four bars) values and predicted values for each of the four models. Filled bars: *t-*test*, p* < 0.05. D, Covariation of *me* (top) and *z* (bottom) with the orientation of the >97% RTrial heatmap patch for monkeys and predictions of the four models. Blue: data from LR-Left blocks, red: data from LR-Right blocks. Data in the “Monkey” column are the same as in Fig 6C and D. Note that predictions of the “over-*me*” model best matched the monkey data than the other models.

The “over-*me*” model produced better predictions than the other three alternative models for all three monkeys. Of the four models, only the “over-*me*” model captured the monkeys’ tendency to bias *me* toward the large-reward choice (positive Δ*me*) and bias *z* toward the small-reward choice (negative Δ*z*; Fig. 8B). In contrast, the “over-*z*” model predicted small adjustments in *me* and large adjustments in *z* favoring the large-reward choice; the “over-both” model predicted relatively large, symmetric *me* and z adjustments favoring the large-reward choice; and the “neutral” model predicted relatively small, symmetric adjustments in both *me* and *z* favoring the large-reward choice. Accordingly, for each monkey, the predicted and actual values of both Δ*me* and Δ*z* were most strongly positively correlated for predictions from the “over-*me*” model compared to the other models (Fig. 8C). The “over-*me”* model was also the only one of the models we tested that recapitulated the measured relationships between both *me*- and *z*-dependent biases and session-by-session changes in the orientation of the RTrial function (Fig. 8D). Qualitatively similar results were obtained with a different criterion that stopped updating when RTrial reached a fixed 98% of RTrial_max_, suggesting that the ability of the “over-*me*” model to mimic monkeys’ performance did not simply result from setting session-specific final RTrial values. Similar results were observed using RR function (Supplementary Figs. 10 and 11).

**Figure S10.**
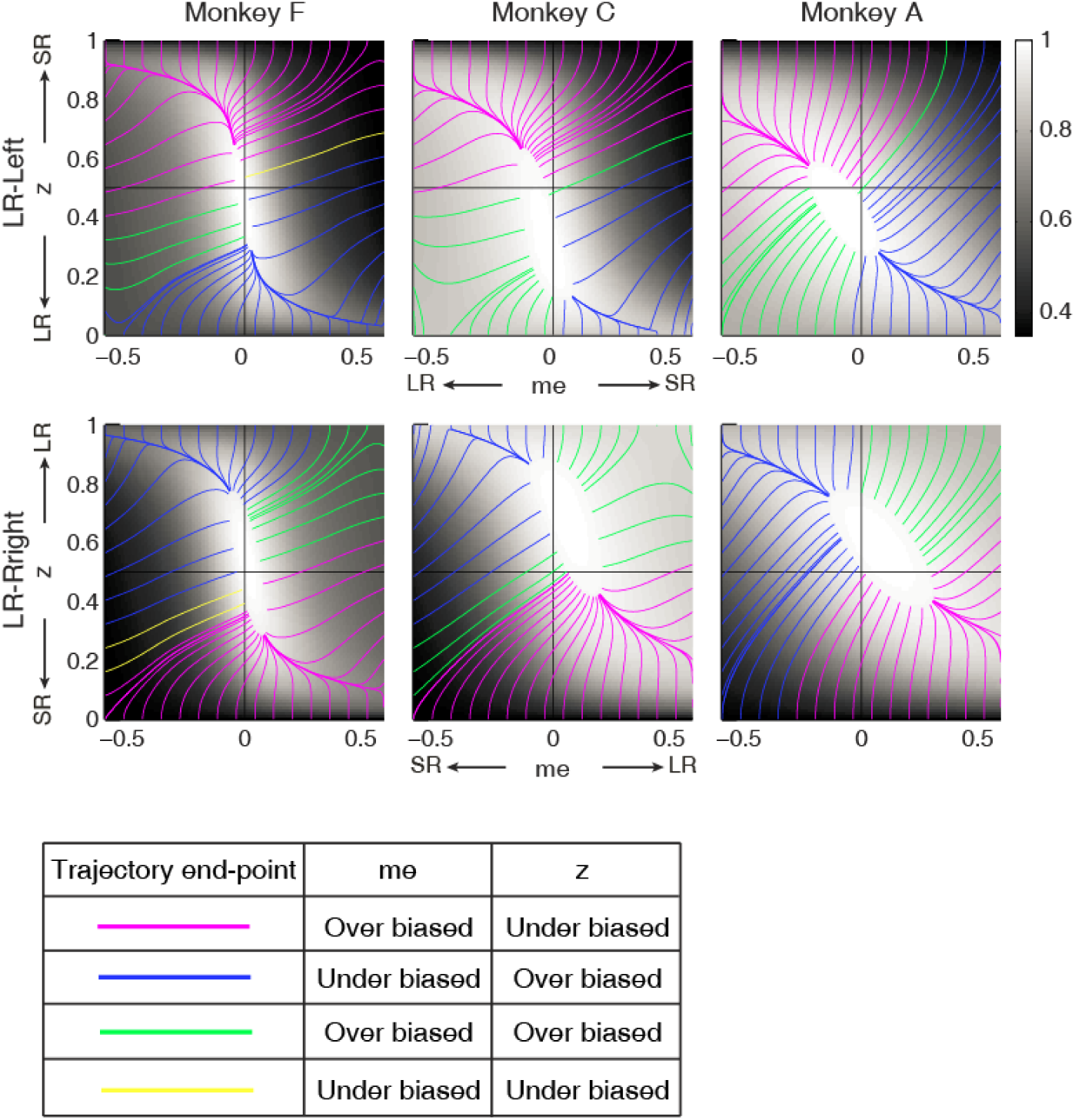
RR gradient trajectories color-coded by the end points of the *me/z* patterns. Same format as Fig. 7 but using gradients based on RR instead of RTrial.

**Figure S11.**
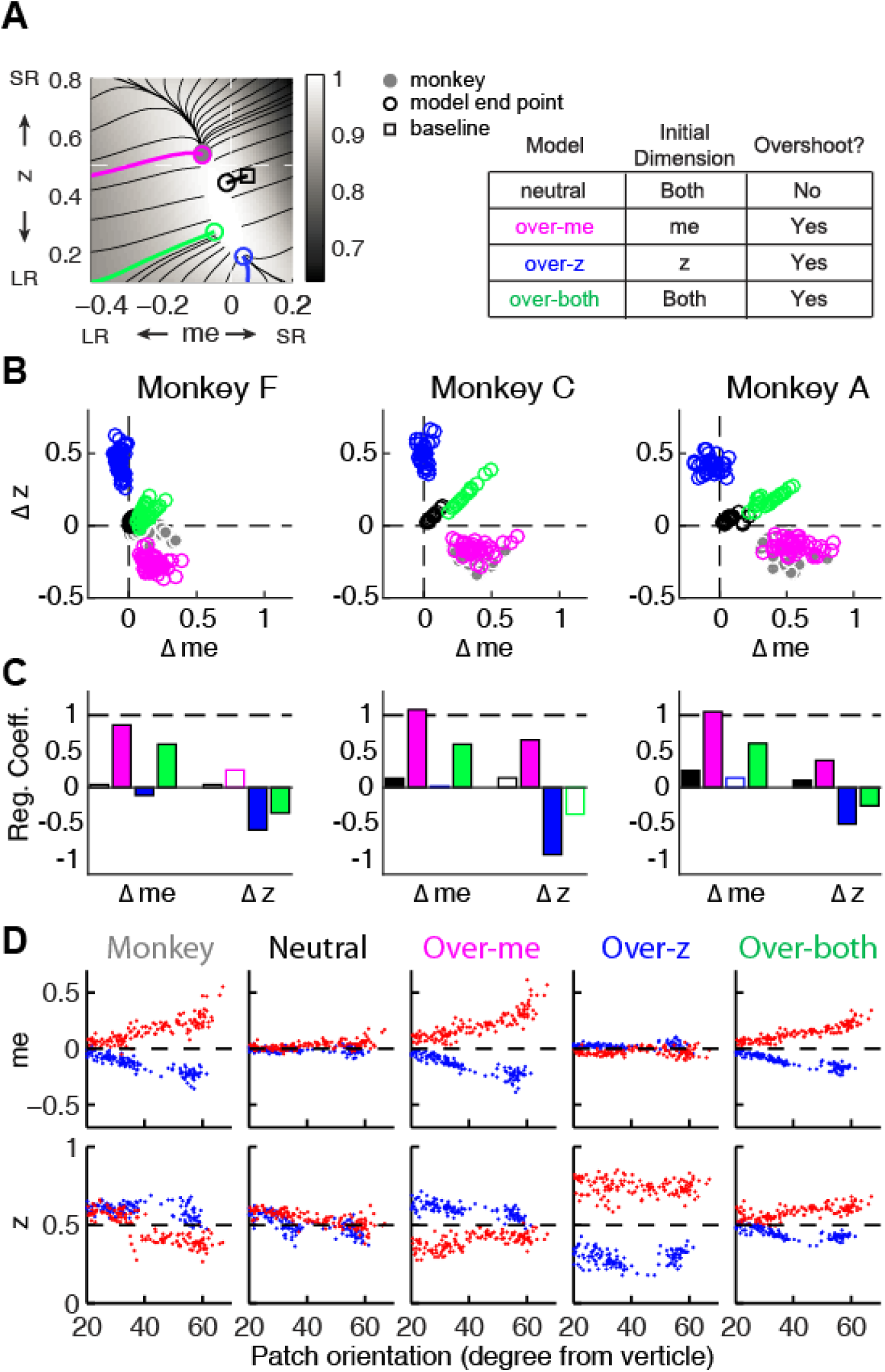
Predictions of a RR gradient-based model. Same format as Fig. 8 but using gradients based on RR instead of RTrial.

## Discussion

We analyzed the behavior of three monkeys performing a decision task that encouraged the use of both uncertain visual motion evidence and the reward context. All three monkeys made choices that were sensitive to the strength of the sensory evidence and were biased toward the larger-reward choice, which is roughly consistent with previous studies of humans and monkeys performing similar tasks (Maddox and Bohil, 1998; Voss et al., 2004; Diederich and Busemeyer, 2006; Liston and Stone, 2008; Serences, 2008; Feng et al., 2009; Simen et al., 2009; Nomoto et al., 2010; Summerfield and Koechlin, 2010; Teichert and Ferrera, 2010; Gao et al., 2011; Leite and Ratcliff, 2011; Mulder et al., 2012; Wang et al., 2013; White and Poldrack, 2014).However, we also found that these adjustments differed considerably in detail for the three monkeys, in terms of overall magnitude, dependence on perceptual sensitivity and offered rewards, and relationship to RTs. We quantified these effects with a logistic analysis and a commonly used model of decision-making, the drift-diffusion model (DDM), which allowed us to compare the underlying decision-related computations to hypothetical benchmarks that would maximize reward. We found that all three monkeys made reward context-dependent adjustments with two basic components: 1) an over-adjustment of the momentary evidence provided by the sensory stimulus (*me*) in favor of the large-reward option; and 2) an adjustment to the decision rule that governs the total evidence needed for each choice (*z*), but in the opposite direction (i.e., towards the small-reward option). These adjustments tended to provide nearly, but not exactly, maximal reward intake. We proposed a common heuristic strategy based on the monkeys’ individual reward functions to account for the idiosyncratic adjustments across monkeys and across sessions within the same monkey.

### Considerations for assessing optimality and rationality

Assessing decision optimality requires a model of the underlying computations. In this study, we chose the DDM for several reasons. First, it provided a parsimonious account of both the choice and RT data (Palmer et al., 2005; Ratcliff et al., 1999). Second, as discussed in more detail below, the DDM and related accumulate-to-bound models have provided useful guidance for identifying neural substrates of the decision process (Roitman and Shadlen, 2002; Ding and Gold, 2010; Ding and Gold, 2012; Hanks et al., 2011; Ratcliff et al., 2003; Rorie et al., 2010; Mulder et al., 2012; Summerfield and Koechlin, 2010; Frank et al., 2015). Third, these models are closely linked to normative theory, including under certain assumptions matching the statistical procedure known as the sequential probability ratio test that can optimally balance the speed and accuracy of uncertain decisions (Barnard, 1946; Wald, 1947; Wald and Wolfowitz, 1948). These normative links were central to our ability to use the DDM to relate the monkeys’ behavior to different forms of reward optimization.

Assessing optimality also requires an appropriate definition of the optimization goal. In our study, we mainly focused on the goal of maximizing reward rate (per trial or per unit of time). Based on this definition, the monkeys showed suboptimal reward-context-dependent adjustments. It is possible that the monkeys’ were optimizing for a different goal, such as accuracy or a competition between reward and accuracy (“COBRA,” Maddox and Bohil, 1998). However, the monkeys’ behavior was not consistent with optimizing for these goals, either. Specifically, none of these goals would predict optimal *z* adjustment that favors the small reward choice: accuracy maximization would require unbiased decisions (*me*=0 and *z*=0.5), whereas COBRA would require *z* values with smaller magnitude (between 0.5 and those predicted for reward maximization alone), but still in the adaptive direction. Therefore, the monkeys’ strategies were not consistent with simply maximizing commonly considered reward functions.

Deviations from optimal behavior are often ascribed to a lack of effort or poor learning. However, these explanations seem unlikely to be primary sources of suboptimality in our study. For example, lapse rates, representing the overall ability to attend to and perform the task, were consistently near zero for all three monkeys. Moreover, the monkeys’ reward outcomes (RTrial or RR with respect to optimal values) did not change systematically with experience but instead stayed close to the optimal values. These results imply that the monkeys understood the task demands and performed consistently well over the course of our study. More importantly, the monkeys made adjustments that were adapted to changes in their idiosyncratic, context-dependent reward functions, which reflected session-specific reward ratios and motion coherences and the monkeys’ daily variations of perceptual sensitivity and speed-accuracy trade-offs (Fig. 6, Supplementary Fig. 7). Based on these observations, we reasoned that the seemingly sub-optimal behaviors may instead reflect a common, adaptive, rational strategy that aimed to attain good-enough (satisficing) outcomes.

The gradient-based, satisficing model we proposed was based on the considerations discussed below to account for our results. We do not yet know how well this model generalizes to other tasks and conditions, but it exemplifies an additional set of general principles for assessing the rationality of decision-making behavior: goals that are not necessarily optimal but good enough potential heuristic strategies based on the properties of the utility function, and flexible adaptation to changes in the external and internal conditions.

### Assumptions and experimental predictions of the proposed learning strategy

In general, finding rational solutions through trial-and-error or stepwise updates requires a sufficient gradient in the utility function to drive learning (Sutton and Barto, 1998). Our proposed scheme couples a standard gradient-following algorithm with principles that have been used to explain and facilitate decisions with high uncertainties, time pressures, and/or complexity to achieve a satisficing solution (Simon, 1966; Wierzbicki, 1982; Gigerenzer and Goldstein, 1996; Nosofsky and Palmeri, 1997; Goodrich et al., 1998; Sakawa and Yauchi, 2001; Goldstein and Gigerenzer, 2002; Stirling, 2003; Gigerenzer, 2010; Oh et al., 2016). This scheme complements but differs from a previously proposed satisficing strategy to account for human subjects’ suboptimal calibration of the speed-accuracy trade-off via adjustments of the decision bounds of a DDM that favor robust solutions given uncertainties about the inter-trial interval (Zacksenhouse et al., 2010), In contrast, our proposed strategy focuses on reward-biased behaviors for a given speed-accuracy tradeoff and operates on reward per trial, which is, by definition, independent of inter-trial-interval. Our scheme was based on three key assumptions, as follows.

Our first key assumption was that the starting point for gradient following was not the unbiased state (i.e., *me*=0 and *z*=0.5) but an over-biased state. Notably, in many cases the monkeys could have performed as well or better than they did, in terms of optimizing reward rate, by making unbiased decisions. The fact that none did so prompted our assumption that their session-by-session adjustments tended to reduce, not inflate, biases. Specifically, we assumed that the initial experience of the asymmetric reward prompted an over-reaction to bias choices towards the large-reward alternative. In general, such an initial over-reaction is not uncommon, as other studies have shown excessive, initial biases that are reduced or eliminated with training (Gold et al., 2008; Jones, et al., 2015; Nikolaev et al., 2016). The over-reaction is also rational because the penalty is larger for an under-reaction than for an over-reaction. For example, in the average RTrial heatmaps for our task (Fig. 6A), the gradient dropped faster in the under-biased side than in the over-biased side. This pattern is generally true for tasks with sigmoid-like psychometric functions (for example, the curves in Supplementary Fig. 2). Our model further suggests that the nature of this initial reaction, which may be driven by individually tuned features of the reward function that can remain largely consistent even for equal-reward tasks (Supplementary Fig. 12) and then constrain the end-points of a gradient-based adjustment process (Fig. 8), may help account for the extensive individual variability in biases that has been reported for reward-biased perceptual tasks (Voss et al., 2004; Summerfield and Koechlin, 2010; Leite and Ratcliff, 2011; Cicmil et al., 2015)).

**Figure S12.**
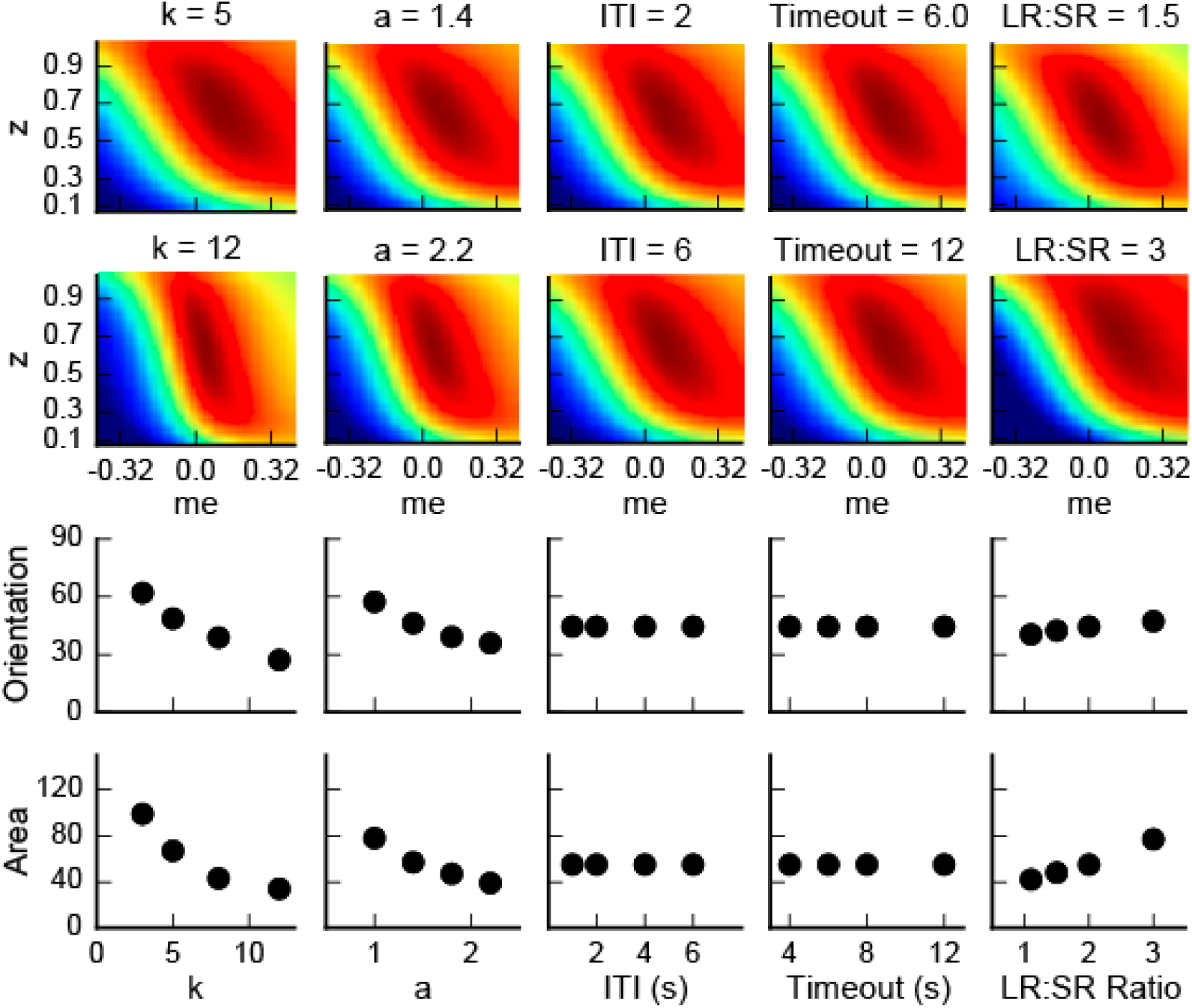
Dependence of the orientation and area of the near-optimal RTrial patch on parameters reflecting internal decision process and external task specifications. The top two rows show the RTrial heatmaps with two values of a single parameter indicated above, while keeping the other parameters fixed at the baseline values. The third and fourth rows show the estimated orientation (the amount of tilt from vertical, in degrees) and area (in pixels), respectively, of the image patches corresponding to ≥97% of RTrial_max_. The baseline values of the parameters are: *a*=1.5, *k*=6, non-decision times=0.3 sec for both choices, *ITI*=4 sec, *Timeout*=8 sec, *large-reward (LR): small-reward (SR) ratio=*2.

The specific form of initial over-reaction in our model, which was based on the gradient asymmetry of the reward function, makes testable predictions. Specifically, our data were most consistent with an initial bias in momentary evidence (*me*), which caused the biggest change in the reward function. However, this gradient asymmetry can change dramatically under different conditions. For example, changes in the subject’s cautiousness (i.e., the total bound height parameter, *a*) and perceptual sensitivity (*k*) would result in a steeper gradient in the other dimension (the decision rule, or *z*) of the reward function (Supplementary Fig. 13). Our model predicts that such a subject would be more prone to an initial bias along that dimension. This prediction can be tested by using speed-accuracy instructions to affect the bound height and different stimulus parameters to change perceptual sensitivity (Palmer et al 2005; Gegenfurtner and Hawken, 1996).

**Figure S13:**
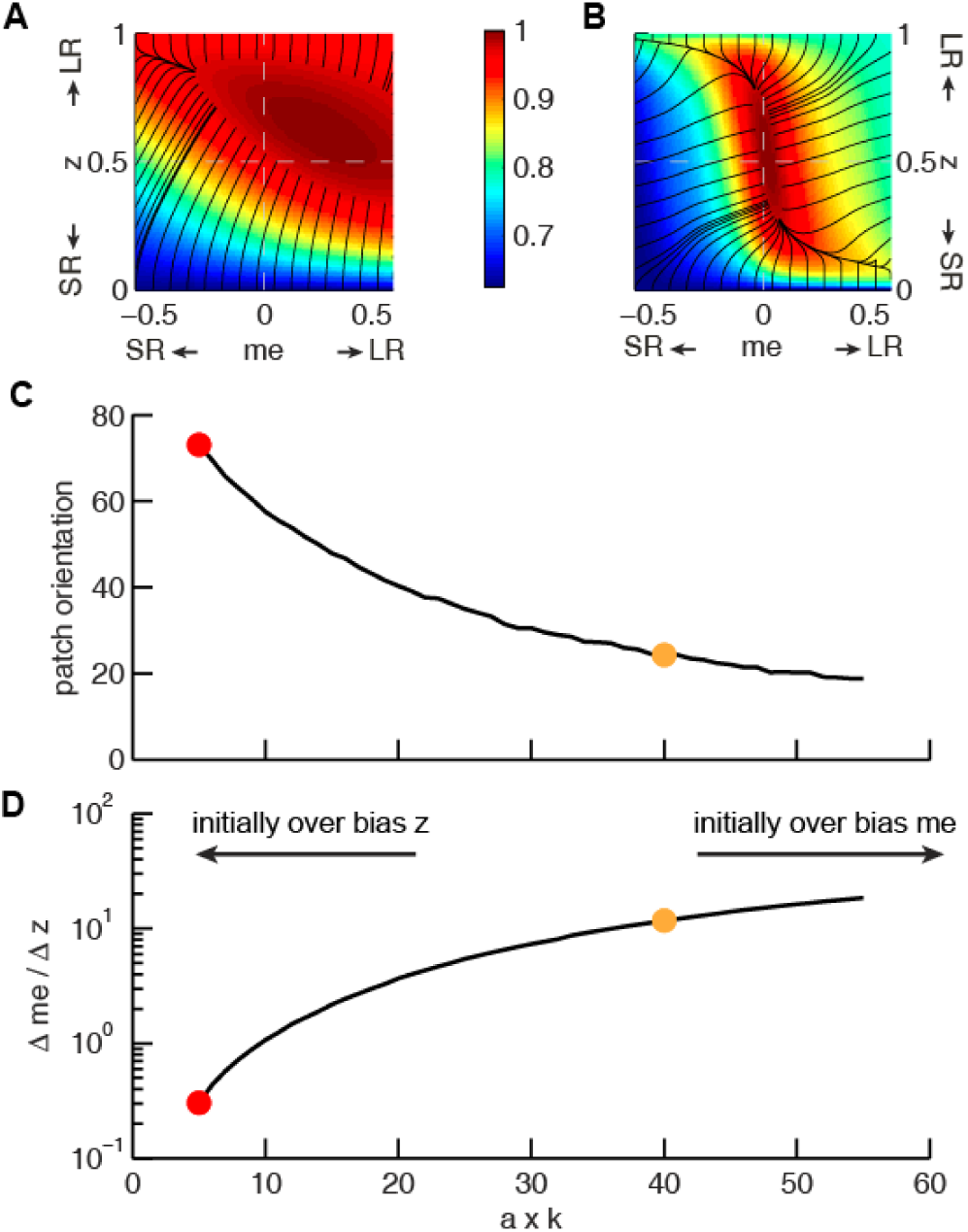
The joint effect of DDM model parameters *a* (governing the speed-accuracy trade-off) and *k* (governing perceptual sensitivity) on the shape of the reward function. A and B, Example RTrial functions corresponding to steeper gradients along the *z* (panel A, corresponding to the red points in panels C and D) or *me* (panel B, corresponding to the orange points in panels C and D) dimension. The gradient lines (black) stop when RTrial >0.97 of the maximum value. A: *a*=1, *k*=5. B: *a*=1, *k*=40. *Large-reward:small-reward ratio* = 2. C, Orientation of the patch corresponding to >0.97 maximal RTrial as a function of the product of *a* and *k*. D, The ratio of the mean gradients along the *me* and *z* dimensions as a function of the product of *a* and *k*. Our model assumes that the initial bias is along the dimension with the steeper gradient according to each monkey’s idiosyncratic RTrial function. Note that because *me* and *z* have different units, the boundary between initial-*me* and initial-*z* conditions may not correspond to a gradient ratio of 1.

Our second key assumption was that from this initial, over-biased state, the monkeys made adjustments to both the momentary evidence (*me*) and decision rule (*z*) that generally followed the gradient of the reward function. The proposed step-wise adjustments occurred too quickly to be evident in behavior; e.g., the estimated biases were similar for the early and late halves in a block (data not shown). Instead, our primary support for this scheme was that the steady-state biases measured in each session were tightly coupled to the shape of the reward function for that session. It would be interesting to design tasks that might allow for more direct measurements of the updating process itself, for example, by manipulating both the initial biases and relevant reward gradient that might promote a longer adjustment process.

Our third key assumption was that the shallowness of the utility of the function around the peak supported satisficing solutions. Specifically, gradient-based adjustments, particularly those that use rapid updates based on implicit knowledge of the utility function, may be sensitive only to relatively large gradients. For our task, the gradients were much smaller around the peak, implying that there were large ranges of parameter values that provided such similar outcomes that further adjustments were not used. In principle, it is possible to change the task conditions to test if and how subjects might optimize with respect to steeper functions around the peak. For example, for RTrial, the most effective way to increase the gradient magnitude near the peak (i.e., reducing the area of the dark red patch) is to increase sensory sensitivity (*k*) or cautiousness (*a*; i.e., emphasizing accuracy over speed; Supplementary Fig. 12). For RR, the gradient can also be enhanced by increasing the time-out penalty. Despite some practical concerns about these manipulations (e.g., increasing time-out penalties can decrease motivation), it would be interesting to study their effects on performance in more detail to understand the conditions under which satisficing or “good enough” strategies are used (Simon, 1956; Simon, 1982).

The satisficing reward gradient-based scheme we propose may further inform appropriate task designs for future studies. For example, our scheme implies that the shape of the reward function near the peak, particularly the steepness of the gradient, can have a strong impact on how closely a subject comes to the optimal solution for a given set of conditions. Thus, task manipulations that affect the shape of the reward-function peak could, in principle, be used to control whether a study focuses on more- or less-optimal behaviors (Supplementary Fig. 14). For example, increasing perceptual sensitivity (e.g., via training) and/or decisions that emphasize accuracy over speed (e.g., via instructions) tends to sharpen the peak of the reward function. According to our scheme, this sharpening should promote increasingly optimal decision-making, above and beyond the performance gains associated with increasing accuracy, because the gradient can be followed closer to the peak of the reward function. The shape of the peak is also affected by the reward ratio, such that higher ratios lead to larger plateaus, i.e. shallower gradient, near the peak. This relationship leads to the idea that, all else being equal, a smaller reward ratio may be more suitable for investigating principles of near-optimal behavior, whereas a larger reward ratio may be more suitable for investigating the source and principles of sub-optimal behaviors.

**Figure S14.**
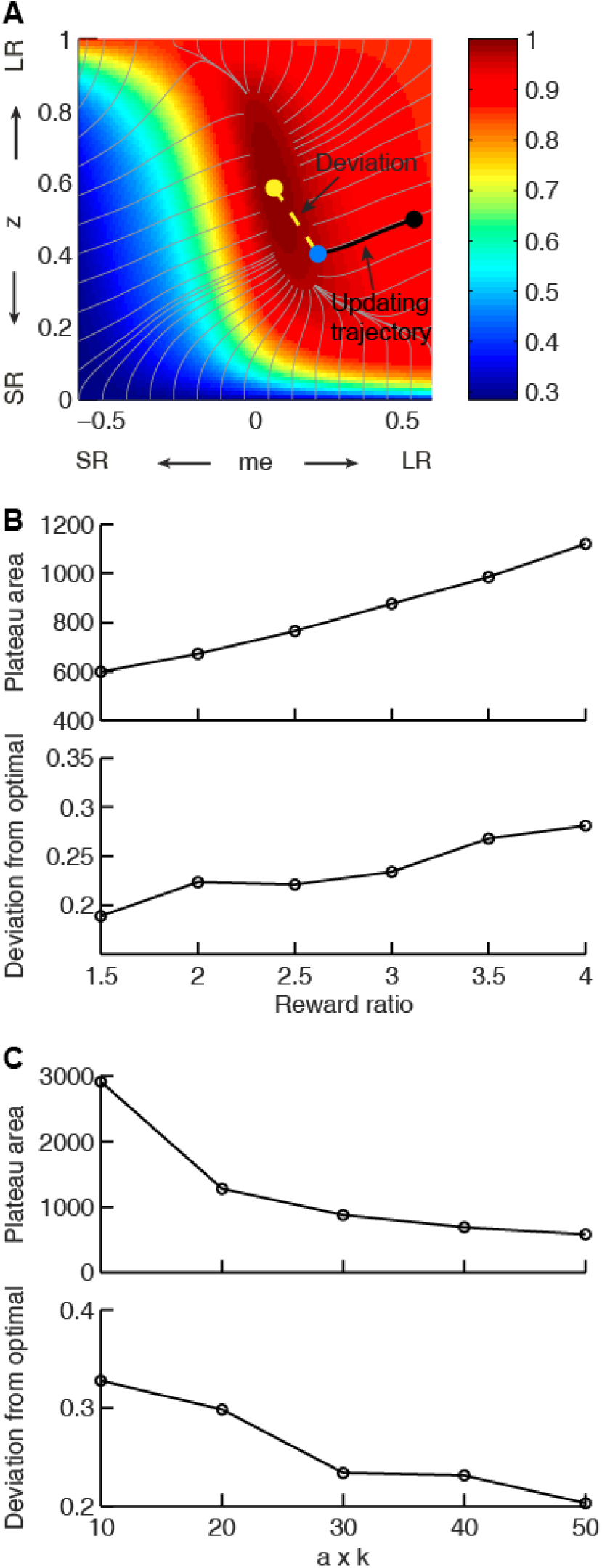
Effects of the shape of the reward function on deviations from optimality. A, Illustration of our heuristic updating model and measurement of deviation of the end point from optimal. Yellow dot: optimal solution. Gray lines: trajectory for gradient ascent, ending at 0.97 maximal RTrial. Black line: trajectory for updating from the starting point (black dot, *me*=0.54, *z*=0.5), which ended at 0.97 maximal RTrial (blue dot). The deviation of the end point from optimal is measured as the distance from the yellow dot to the blue dot (yellow dashed line). The same starting point and ending criterion were used for data shown in B and C. B, The area of the 0.97 maximal RTrial plateau and end-point deviation from optimal increase with reward ratio. The product of *a* and *k* is fixed as 30. C, The area of the 0.97 maximal RTrial plateau and end-point deviation from optimal decrease with the product of *a* and *k*. Reward ratio is fixed as 3.

### Possible neural mechanisms

The DDM framework has been used effectively to identify and interpret neural substrates of key computational components of the decision process for symmetric-reward versions of the motion-discrimination task. Our study benefitted from an RT task design that provided a richer set of constraints for inferring characteristics of the underlying decision process than choice data alone (Feng et al., 2009; Nomoto et al., 2010; Teichert and Ferrera, 2010). The monkeys’ strategy further provides valuable anchors for future studies of the neural mechanisms underlying decisions that are biased by reward asymmetry, stimulus probability asymmetry, and other task contexts.

For neural correlates of bias terms in the DDM, it is commonly hypothesized that *me* adjustments may be implemented as modulation of MT output and/or synaptic weights for the connections between different MT subpopulations and decision areas (Cicmil, et al., 2015). In contrast, *z* adjustments may be implemented as context-dependent baseline changes in neural representations of the decision variable and/or context-dependent changes in the rule that determines the final choice (Lo and Wang, 2006; Rao, 2010; Lo et al., 2015; Wei et al., 2015). The manifestation of these adjustments in neural activity that encodes a decision variable may thus differ in its temporal characteristics: a *me* adjustment is assumed to modulate the rate of change in neural activity, whereas a *z* adjustment does not. However, such a theoretical difference can be challenging to observe, because of the stochasticity in spike generation and, given such stochasticity, practical difficulties in obtaining sufficient data with long decision deliberation times. By adjusting *me* and *z* in opposite directions, our monkeys’ strategies may allow a simpler test to disambiguate neural correlates of *me* and *z*. Specifically, a neuron that encodes *me* may show reward modulation congruent with its choice preference, whereas a neuron that encodes *z* may show reward modulation opposite to its choice preference (Supplementary Fig. 15). These predictions further suggest that, although it is important to understand if and how human or animal subjects can perform a certain task optimally, for certain systems-level questions, there may be benefits to tailoring task designs to promote sub-optimal strategies in otherwise well-trained subjects.

**Figure S15.**
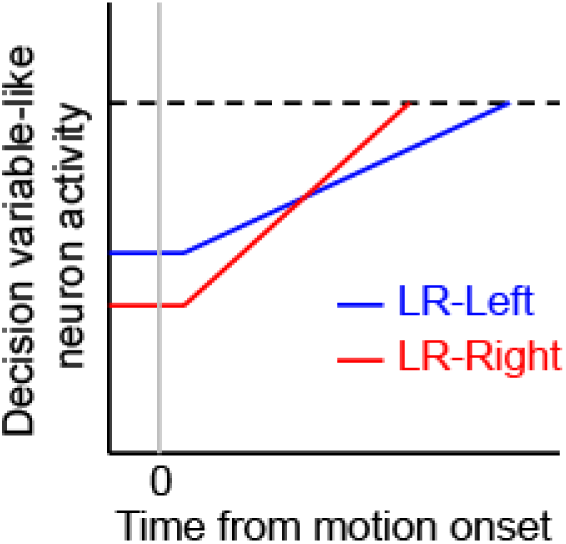
Hypothetical neural activity encoding a reward-biased perceptual decision variable. The blue and red curves depict rise-to-threshold dynamics in favor of a particular (say, rightward) choice under the two reward contexts, as indicated. Note that when the rightward choice is paired with larger reward: 1) the slope of the ramping process, which corresponds to an adjustment in momentary evidence (*me*), is steeper; and 2) the baseline activity, which corresponds to the decision-rule (*z*) adjustment, is lower.

## Methods

### Subjects

We used three rhesus macaques (*Macaca mulatta*), two male and one female, to study behavior on an asymmetric-reward reaction-time random-dot motion discrimination task (Fig. 1B, see below). Prior to this study, monkeys F and C had been trained extensively on the equal-reward RT version of the task (Ding and Gold, 2010, 2012b, a). Monkey A had been trained extensively on non-RT dots tasks (Connolly et al., 2009; Bennur and Gold, 2011), followed by >130 sessions of training on the equal-reward RT dots task. All training and experimental procedures were in accordance with the National Institutes of Health Guide for the Care and Use of Laboratory Animals and were approved by the University of Pennsylvania Institutional Animal Care and Use Committee.

### Behavioral task

Our task (Fig. 1B) was based on the widely used random-dot motion discrimination task that typically has symmetric rewards (Roitman and Shadlen, 2002; Ding and Gold, 2010). Briefly, a trial started with presentation of a fixation point at the center of a computer screen in front of a monkey. Two choice targets appeared 0.5 s after the monkey acquired fixation. After a delay, the fixation point was dimmed and a random-dot kinematogram (speed: 6 °/s) was shown in a 5° aperture centered on the fixation point. For monkeys F and C, the delay duration was drawn from a truncated exponential distribution with mean=0.7 s, max=2.5 s, min=0.4 s. For monkey A, the delay was set as 0.75 s. The monkey was required to report the perceived global motion direction by making a saccade to the corresponding choice target at a self-determined time (a 50-ms minimum latency was imposed to discourage fast guesses). The stimulus was immediately turned off when the monkeys’ gaze left the fixation window (4, 4, and 3° square windows for monkey F, C, and A, respectively). Correct choices (i.e., saccades to the target congruent with actual motion direction) were rewarded with juice. Error choices were not rewarded and instead penalized with a timeout before the next trial began (timeout duration: 3 s, 0.5-2 s, and 2.5 s, for monkeys F, C, and A, respectively).

On each trial, the motion direction was randomly selected toward one of the choice targets along the horizontal axis. The motion strength of the kinematogram was controlled as the fraction of dots moving coherently to one direction (coherence). On each trial, coherence was randomly selected from 0.032, 0.064, 0.128, 0.256, and 0.512 for monkeys F and C, and from 0.128, 0.256, 0.512, and 0.75 for monkey A. In a subset of sessions, coherence levels of 0.064, 0.09, 0.35, and/or 0.6 were also used for monkey A.

We imposed two types of reward context on the basic task. For the “LR-Left” reward context, correct leftward saccades were rewarded with a larger amount of juice than correct rightward saccades. For the “LR-Right” reward context, correct leftward saccades were rewarded with a smaller amount of juice than correct rightward saccades. The large:small reward ratio was on average 1.34, 1.91, and 2.45 for monkeys F, C, and A, respectively. Reward context was alternated between blocks and constant within a block. Block changes were signaled to the monkey with an inter-block interval of 5 s. The reward context for the current block was signaled to the monkey in two ways: 1) in the first trial after a block change, the two choice targets were presented in blue and green colors, for small and large rewards, respectively (this trial was not included for analysis); and 2) only the highest coherence level (near 100% accuracy) was used for the first two trials after a block change to ensure that the monkey physically experienced the difference in reward outcome for the two choices. For the rest of the block, choice targets were presented in the same color and motion directions and coherence levels were randomly interleaved.

### Basic characterization of behavioral performance

Eye position was monitored using a video-based system (ASL) sampled at 240 Hz. RT was measured as the time from stimulus onset to saccade onset, the latter identified offline with respect to velocity (> 40°/s) and acceleration (> 8000°/s^2^). Performance was quantified with psychometric and chronometric functions (Figs. 2 and 3), which describe the relationship of motion strength (signed coherence, *Coh*, which was the proportion of the dots moving in the same direction, positive for rightward motion, negative for leftward motion) with choice and RT, respectively. Psychometric functions were fitted to a logistic function (Equation (1)), in which λ is the error rate, or lapse rate, independent of the motion information; α_0_ and (α_0_ + α_rew_) are the bias terms, which measures the coherence at which the performance was at chance level in the LR-Right and LR-Left reward contexts, respectively. β_0_ and (β_0_ + β_rew_) are the perceptual sensitivities in the LR-Right and LR-Left reward contexts, respectively.

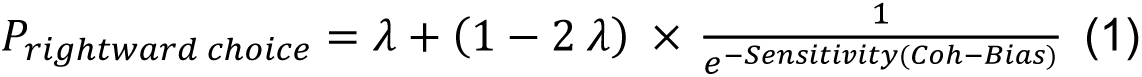

### Reward-biased drift-diffusion model

To infer the computational strategies employed by the monkeys, we adopted the widely used accumulation-to-bound framework, the drift-diffusion model (DDM; Fig. 1A). In the standard DDM, motion evidence is modeled as a random variable following a Gaussian distribution with a mean linearly proportional to the signed coherence and a fixed variance. The decision variable (DV) is modeled as temporal accumulation (integral) of the evidence, drifting between two decision bounds. Once the DV crosses a bound, evidence accumulation is terminated, the identity of the decision is determined by which bound is crossed, and the decision time is determined by the accumulation time. RT is modeled as the sum of decision time and saccade-specific non-decision times, the latter accounting for the contributions of evidence-independent sensory and motor processes.

To model the observed influences of motion stimulus and reward context on monkeys’ choice and RT behavior, we introduced two reward context-dependent terms: *z* specifies the relative bound heights for the two choices and *me* specifies the equivalent momentary evidence that is added to the motion evidence at each accumulating step. Thus, for each reward context, six parameters were used to specify the decision performance: *a:* total bound height; *k*: proportional scaling factor converting evidence to the drift rate; *t_0_* and *t_1_*: non-decision times for leftward and rightward choices, respectively; and *z* and *me.* Similar approaches have been used in studies of human and animal decision making under unequal payoff structure and/or prior probabilities (Voss et al., 2004; Bogacz et al., 2006; Diederich and Busemeyer, 2006; Summerfield and Koechlin, 2010; Hanks et al., 2011; Mulder et al., 2012).

To fit the monkeys’ data, we implemented hierarchical DDM fitting using an open-source package in Python, which performs Bayesian estimates of DDM parameters based on single-trial RTs (Wiecki et al., 2013). This method assumes that parameters from individual sessions are samples from a group distribution. The initial prior distribution of a given parameter is determined from previous reports of human perceptual performance and is generally consistent with monkey performance on equal reward motion discrimination tasks (Ding and Gold, 2010; Matzke and Wagenmakers, 2009). The posterior distributions of the session- and group-level parameters are estimated with Markov chain Monte Carlo sampling.

For each dataset, we performed 5 chains of sampling with a minimum of 10000 total samples (range: 10000-20000; burn-in: 5000 samples) and inspected the trace, autocorrelation and marginal posterior histogram of the group-level parameters to detect signs of poor convergence. To ensure similar level of convergence across models, we computed the Gelman-Rubin statistic (R-hat) and only accepted fits with R-hat<1.01.

To assess whether reward context modulation of both *z* and *me* was necessary to account for monkeys’ behavioral data, we compared fitting performance between the model with both terms (“full”) and reduced models with only one term (“z-only” and “me-only”). Model selection was based on the deviance information criterion (DIC), with a smaller DIC value indicating a preferred model. Because DIC tends to favor more complex models, we bootstrapped the expected ΔDIC values, assuming the reduced models were the ground truth, using trial-matched simulations. For each session, we generated simulated data using the DDM, with single-session parameters fitted by *me*-only or *z*-only HDDM models and with the number of trials for each direction × coherence × reward context combination matched to the monkey’s data for that session. These simulated data were then re-fitted by all three models to estimate the predicted ΔDIC, assuming the reduced model as the generative model.

To test an alternative model, we also fitted monkeys’ data to a DDM with collapsing bounds (Zylberberg et al., 2016). This DDM was constructed as the expected first-stopping-time distribution given a set of parameters, using the PyMC module (version 2.3.6) in Python (version 3.5.2). The three model variants, “full”, “*me*-only” and “*z*-only”, and their associated parameters were the same as in HDDM, except that the total bound distance decreases with time. The distance between the two choice bounds is set as *a*/(1 + *e*^*β*(*t*−*d*)^), where *a* is the initial bound distance, *β* determines the rate of collapsing, and *d* determines the onset of the collapse. Fitting was performed by computing the maximum *a posteriori* estimates, followed by Markov chain Monte Carlo sampling, of DDM parameters given the experimental RT data.

### Optimality analysis

To examine the level of optimality of the monkeys’ performance, we focused on two reward functions: reward rate (RR, defined as the average reward per second) and reward per trial (RTrial, defined as the average reward per trial) for a given reward context for each session. To estimate the reward functions in relation to *me* and *z* adjustments for a given reward context, we numerically obtained choice and RT values for different combinations of *z* (ranging from 0 to 1) and *me* (ranging from -0.6 to 0.6 coherence unless otherwise specified), given *a, k* and non-decision time values fitted by the full model. We then calculated RR and RTrial, using trial-matched parameters, including the actual ITI, timeout, and large:small reward ratio. RR_max_ and RTrial_max_ were identified as the maximal values given the sampled *me-z* combinations, using 1000 trials for each coherence × direction condition. Optimal *me* and *z* adjustments were defined as the *me* and *z* values corresponding to RR_max_ or RTrial_max_. RR_predict_ and RTrial_predict_ were calculated with the fitted *me* and *z* values in the full model.

## Acknowledgments

We thank Takahiro Doi for helpful comments, Javier Caballero and Rachel Gates for animal training, Jean Zweigle for animal care, and Michael Yoder for data entry.

